# Metallo-protease Peptidase M84 from *Bacillus altitudinis* induces ROS dependent apoptosis in ovarian cancer cells by targeting PAR-1

**DOI:** 10.1101/2023.09.06.556500

**Authors:** Niraj Nag, Tanusree Ray, Rima Tapader, Animesh Gope, Rajdeep Das, Elizabeth Mahapatra, Saibal Saha, Ananda Pal, Parash Prasad, Shruti Chatterjee, Sib Sankar Roy, Amit Pal

## Abstract

In pursuit of isolating novel anticancer proteases from environmental microbial isolates, we have purified and identified an extracellular metallo-protease from *Bacillus altitudinis* named Peptidase M84. This protease selectively triggered apoptosis in human ovarian adenocarcinoma cells (PA-1, SKOV3) and mouse ovarian carcinoma cells (ID8), in addition to exhibiting no significant effect on normal human epithelial ovarian cell (IOSE) and mouse peritoneal macrophage (PEMФ) cell viabilities. Protease activated receptor-1 (PAR-1); a GPCR which is reported to be overexpressed in ovarian cancer cells was identified as a novel target of Peptidase M84. We observed that Peptidase M84 induced PAR-1 overexpression along with activating its downstream signalling effectors NFκB and MAPK to promote excessive reactive oxygen species (ROS) generation in ovarian cancer cells. This disrupted mitochondrial membrane potential, allowed cytosolic release of mitochondrial cytochrome c, increased the Bax (pro-apoptotic) to Bcl-2 (anti-apoptotic) ratio and promoted DNA damage to evoke apoptotic death of the ovarian cancer cells. Peptidase M84 also reduced nuclear ki-67 expression in these malignant cells to render an anti-proliferative role. In *in vivo* set-up, weekly intraperitoneal administration of Peptidase M84 (12 µg/kg body-weight) in the ID8 mice model significantly diminished ascitic fluid accumulation through induction of oxidative stress, increasing murine survival rates by 60%. Collectively, our *in vitro* and *in vivo* findings suggested that Peptidase M84 triggered PAR-1 mediated oxidative stress to act as an apoptosis inducer in ovarian cancer cells. This established Peptidase M84 as a promising drug candidate for receptor mediated targeted-therapy of ovarian cancer.

**Graphical Abstract:** 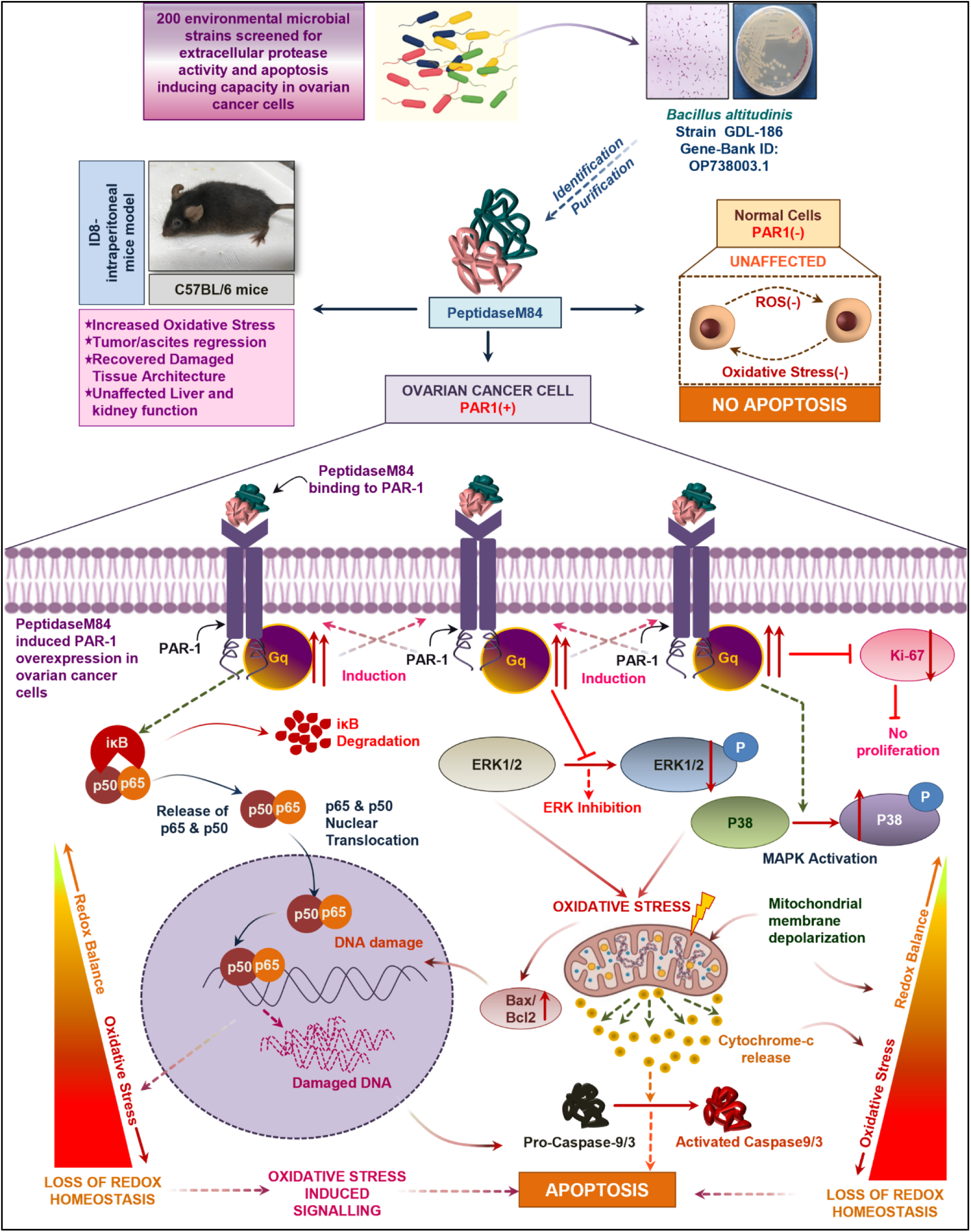

## 1. Introduction

In spite of remarkable improvements in chemotherapy regimens to treat ovarian cancer, it still remains one of the most lethal gynaecological malignancies with a high recurrence and mortality rate [1]. Although cytoreductive surgery accompanied with platinum-based chemotherapy is the admissible treatment for epithelial ovarian cancer (EOC), the mortality rates have not improved markedly [1–3]. For Instance, ovarian cancer is ranked eighth among the top ten common malignancies affecting Indian females as per Globocan 2018. Therefore, new therapies and alternative targets need to be developed to combat ovarian cancer.

In this regard, the anticancer potentials of natural products have been extensively explored for their differential effects as ‘cytoprotectors’ to normal cells and ‘cytotoxics’ to cancer cells. Predominantly, natural products exhibit this dual role by targeting several key molecules which in many instances impart resistance to conventional chemotherapy [4–6]. Herein, microorganisms supply structurally diverse natural products in abundance, serving the purpose of drug discovery. Extraction and Purification of natural bacterial products are relatively cost-effective compared to chemically synthesised drugs [7, 8]. Bacterial toxins and enzymes have been acknowledged for their anti-cancer effects. Reports are suggestive of their role as modulators of various cellular processes like apoptosis, differentiation and proliferation. Bacterial products alone or when conjugated with other available anticancer drugs or irradiation can improve the efficiency of cancer therapy [9–11]. Didemnin B from *Trididemnum solidum* was among the earliest anti-cancer drugs of marine origin to enter clinical studies [12]. Similarly, Azurin from *Pseudomonas aeruginosa* and MakA from *Vibrio cholerae* have already shown promising anticancer effects [13–15]. Microbial proteases are reported to play their role through several mechanisms such as the inactivation of antimicrobial peptides [16], disruption of the defensive mucosal barrier [17] and elicitation of apoptosis in target cells [18]. For example, a protease obtained from *Serratia mercescens kums 3958* was reported to cause significant tumour regression when injected into solid tumours in BALB/c mice. [19, 20]. Previously, we have also reported bacterial subtilisin, a serine protease capable of potentiating apoptosis via ubiquitin mediated tubulin degradation pathway in breast cancer cells [21]. L-asparaginase (ASNase) is a pharmaceutically and clinically important microbial enzyme, isolated from different environmental sources that showed hopeful outcomes in cancer therapy as well [22, 23]. Studies on extracellular metallo-protease arazyme from *Serratia proteamaculans* stated that it could effectively inhibit metastatic murine melanoma via matrix metallo protease-8 cross-reactive antibody stimulation [24]. These findings revealed a possible mechanism of cytotoxic action of microbial proteases on cancer cells.

Oxidative stress serves as a key modulator of signalling pathways involved in cellular survival and death depending on the cellular threshold levels of ROS. Dynamic Redox homeostasis is thus a characteristic of exorbitantly growing cancer cells. Hence, most cytotoxic cancer drugs are observed to promote cell death by inducing oxidative stress, either directly or indirectly [25, 26] Higher levels of cellular ROS persuade mitochondria dependent conventional intrinsic pathway of apoptosis following caspase-9 activation [26, 27]. Proteases can induce bio-signalling pathways and control cellular functions through the cleavage of protease-activated receptors (PARs), a particular G-Protein Coupled Receptors (GPCRs) [28]. The subtypes of PARs, ranging from PAR 1-4, have been found to be overexpressed in cancer cells with respect to healthy normal cells [29]. PARs play important roles in the apoptosis of cancer cells and carcinogenesis depending on the stimuli [30–33]. Studies demonstrated the differential expression pattern of PAR-1 in both transcriptional and translational levels in ovarian carcinoma tissue samples, while, negligible or none in the normal ovarian surface epithelium [34]. Moreover, PAR-1 agonists also showed apoptosis in intestinal epithelial cells [35]. Nevertheless, the molecular mechanism by which overexpression of PAR-1 can perturb the viability of cancer cells remains a relatively less examined area of cancer research. PAR-1 activation can trigger ROS generation in cancer cells [36, 37]. However, both PAR-1 and ROS are also reported to be associated with the activation of NFκB and MAP kinases imparting apoptosis of cancer cells [37, 38]. Therefore, PAR-1 mediated ROS targeting therapy using proteases may be proven to be advantageous to selectively kill cancer cells.

The objective of our study, therefore involved screening of environmental microbial isolates to identify a naturally occurring microbial protease with apoptosis capacitating potentials and deciphering its anticancer mechanism. In this study, we report that Peptidase M84, an extracellular metallo-protease purified from an environmental isolate of gram-positive *Bacillus altitudinis* can augment oxidative stress by altering PAR-1 activity to trigger apoptotic signalling in human and mice ovarian carcinoma cells with high selective toxicity. Our present study provides a deeper insight into the mechanism underlying Peptidase M84 induced apoptosis in ovarian cancer cells in a PAR-1/ROS-dependent manner which was not elucidated previously. Thus, Peptidase M84 may be employed for its chemotherapeutic efficacy in the future for ovarian cancer amelioration.

## 2. Material and Methods

### 2.1 Chemicals and reagents

Different biochemical assay kits (AST, ALT, Urea, Creatinine), JC-1 staining powder (Invitrogen™, Cat# T3168), 2′,7′-Dichlorodihydrofluorescein diacetate (Sigma, Cat# D6883), Mito tracker green FM (Invitrogen™, Cat# M7514), ML-161(Sigma, Cat# SML0418), SB203580 (Sigma, Cat# 559389), MG132 (Sigma, Cat# M7449), DAPI (Sigma, Cat# D9542), Hoechst 33342 (Invitrogen™, Cat# H21492) and other general chemicals of analytical grade were purchased from Sigma-Aldrich (USA) and Invitrogen (USA). Foetal bovine serum (FBS) (Gibco, Cat# 10270106) antibiotics (streptomycin, penicillin) (Gibco, Cat # 15140122) and all cell culture media were purchased from Gibco (USA). Anti-cleaved PARP antibody (Cell Signaling Technology Cat# 5625), anti Bax antibody (Cell Signaling Technology Cat# 5023), anti Bcl-2 antibody (Cell Signaling Technology Cat# 15071), anti-caspase 8 antibody (Cell Signaling Technology Cat# 9746), anti-Cytochrome c antibody (Cell Signaling Technology Cat# 12963 and Cell Signaling Technology Cat# 11940), anti-Histone H3 antibody (Cell Signaling Technology Cat# 14269), anti-α-Tubulin antibody (Cell Signaling Technology Cat# 3873), anti GAPDH antibody (Cell Signaling Technology Cat# 97166), anti-Ki-67 antibody (Cell Signaling Technology Cat# 9449), anti-cleaved caspase 9 antibody (Cell Signaling Technology Cat# 7237), anti-p65 antibody (Cell Signaling Technology Cat# 4764) and anti-p38 antibody (Cell Signaling Technology Cat# 8690) were purchased from cell signalling technology (USA). Anti-PAR-1 antibody (Millipore Cat# MABF244) was purchased from Merck (USA). Anti-ERK1/2 antibody (Santa Cruz Biotechnology Cat# sc-514302), anti-phospho-ERK1/2 antibody (Santa Cruz Biotechnology Cat# sc-136521), anti-phospho-p38 antibody (Santa Cruz Biotechnology Cat# sc-166182) and anti-p50 antibody (Santa Cruz Biotechnology Cat# sc-8414) were purchased from Santacruz biotechnology (USA). Anti-cleaved caspase 3 antibody (Abcam Cat# ab13847), was purchased from Abcam (UK). All secondary antibodies with different conjugates (HRP, AP and Alexa fluor) were purchased from cell signalling technology (USA). All primers (PAR-1, PAR-2, PAR-3, PAR-4, bacterial Peptidase M84 and bacterial DNA gyrase B) for PCR were purchased from IDT (Integrated Technology Enterprise Inc, USA). Lipofectamine™ 3000 transfection reagent (Invitrogen^TM^, Cat# L3000001) and human PAR-1 si-RNA (CGGUCUGUUAUGUGUCUAUdTdT) were purchased from Invitrogen (USA) and Eurogentec (Belgium) respectively. Protein A (CST, Cat# 9863S) and G agarose (CST# 37478S) beads were purchased from cell signalling technology (USA). DEAE-52 matrix (Whatman; Cat# 4057050) and sephadex G-75 matrix (MP bio; Cat# 195589) were purchased from Whatman (UK) and MP biomedical (USA) respectively. BCA assay kit (Thermo Scientific™, Cat# 23225) and DNA ladder (Invitrogen™, Cat# 15628050) were purchased from Thermofisher, (USA). Protein ladder was purchase from Abcam, (Cat# ab116028) (UK) and G-bioscience, (Cat# 786419) (USA). All tissue culture plasticwares were purchased from nunc (USA) and corning (USA).

### 2.2 Bacterial isolates and its growth conditions

A total of 200 bacterial isolates were isolated from salt farm of CSIR-CSMCRI, Bhavnagar (21°47’50’’ N, 72°07’16’’ E), Gujrat, India. All the isolates were stored in 25% glycerol at −80°C. Bacterial isolates were revived in nutrient broth (BD, USA; Cat# 234000) of pH 8.0 containing 2% NaCl (BD, USA; Cat# 234000). Initially, isolates were grown in 5.0 ml NB (primary culture) at 37°C in a shaker incubator till the OD600 reached 0.6. Secondary cultures were given in 1000 ml NB at a ratio of 1:100 and grown at 37°C in a shaker incubator for 18 h. The culture supernatant of the bacterial isolates was used for screening of protease.

### 2.3 Azocasein assay

Azocasein assay was performed to determine the protease activity of the bacterial isolates as described earlier (Syngkon *et al*., 2010) [39]. Briefly, the overnight grown culture supernatants were mixed with 0.7% azocasein (prepared in 100 mM Tris-HCl; pH 8.0) (Sigma, Cat# A2765) followed by incubation at 37°C for 1 h. The reaction was stopped using 10% Tri Chloro Acetic Acid (Sigma; Cat# T6399) and kept at 4°C for 30 min. Precipitate was removed by centrifugation (12,000 rpm for 10 min). NaOH (500 mM) was added to the supernatant and absorbance was measured at 440 nm. Nutrient broth and purified haemagglutinin protease (HAP) from *Vibrio cholerae* C6709 were used as negative and positive controls respectively.

### 2.4 Purification and identification of protease

Peptidase M84 was purified from the culture supernatant of ‘GDL-186’. Bacterial strain was grown in 1000 ml NB (containing 2% NaCl, pH 8.0) for 19 h at 37°C in a shaker incubator. After centrifugation at 12,000 rpm for 10 min at 4°C, the culture supernatant was salted out with 80% saturated ammonium sulphate (Sigma; Cat# A4915) and kept overnight (O/N) at 4°C. Protein pellet was collected by centrifugation at 14000 rpm at 4°C and the pellet was dissolved in 25 mM Tris-Cl buffer, pH 8.0. The protein solution was dialysed against the same buffer for 48 h at 4°C using dialysis membrane (Himedia; Cat# LA395-30MT). Dialysed protein solution was concentrated using speed-vac vacuum centrifugation and run on DEAE-52 ion-exchange column (2.5 × 20 cm) pre-equilibrated with 25 mM Tris-Cl buffer, pH 8.0. Flow through or non-binding fraction was collected, concentrated and checked for protease activity. Binding fractions were collected using increasing concentration of NaCl (0.1 M - 0.5 M) solution. The eluted fractions were pooled, dialysed, concentrated and examined for protease activity. Non-binding fraction showing protease activity, was further pooled, concentrated and loaded into Sephadex G-75 gel filtration column (1.5 × 30 cm) with 25 mM Tris-HCl buffer of pH 8.0. Fractions positive for protease activity were eluted, concentrated and analysed by SDS and Native-PAGE. The single band from Native-PAGE and two bands from SDS-PAGE were cut out from the gel and sent to C-CAMP, NCBS, Bangalore, India for identification by nano-LC-MS/MS-TOF. Peptide sequence generated from MS/MS spectra was searched in NCBI and Uniprot databases for homology alignment.

### 2.5 Determination of physico-chemical characteristics of the purified protease

Inhibition of the protease activity with different inhibitors was performed in order to determine the type of the purified protease and the culture supernatant of isolate GDL-186. PMSF (10 mM) (Sigma; Cat# P7626), EDTA (10 mM) (Sigma; Cat# E9884) and 1,10 phenanthroline (10 mM) (Sigma, Cat# 131377) were used in the inhibition study as described in our previous study (Tapader *et al*, 2016) [40]. 100 mM stocks of PMSF and 1,10 phenanthroline were prepared in isopropanol and methanol respectively and 500 mM stock of EDTA was prepared in water. 5.0 µg of purified protease was pre-incubated at 37°C for 1 h with each inhibitor. The residual protease activity was measured by azocasein assay.

The optimum pH for protease activity was determined using buffers of different pH ranging from 4.0–11.0. 5.0 µg of purified protease was dialysed overnight against 25 mM acetate buffer (pH 4.0–5.0), 25 mM phosphate buffer (pH 6.0–7.0), 25 mM Tris-HCl (pH 8.0–9.0), 25 mM glycine-NaOH buffer (pH 10.0–11.0) and activity were determined with azocasein assay.

To determine the optimum temperature for activity, 5.0 µg of the purified protease was incubated over a wide range of temperatures: 4°C, 25°C, 37°C, 50°C, 60°C and 70°C for 1 h and the enzyme activity was determined by azocasein assay as already described.

Gelatine zymography was performed to determine the substrate specificity of the purified protease as per the protocol described earlier (Tapader *et al*, 2016) [40]. For native zymography, samples were electrophoresed under non-reducing conditions without boiling using 7.5% Native-PAGE co-polymerised with 0.1% gelatin (Sigma; Cat# G2500). The gel was incubated after electrophoresis in renaturation buffer (2.5% Triton-X-100) for 1 h at room temperature (RT) with gentle shaking. The gel was then developed in development buffer containing 5 mM CaCl2, 25 mM Tris-HCl (pH 8.0) for O/N at 37°C with gentle shaking. The gel was visualised after staining using Coomassie Brilliant Blue G-250 (Himedia; Cat# ML046) and subsequent destaining.

## 2.6 16s-rRNA sequencing

The protease positive bacterial isolate which showed significant apoptotic activity was identified by 16s rRNA sequencing. First, genomic DNA was isolated from O/N grown bacterial culture and run on 1.0% Agarose gel.16S rRNA gene was amplified by 27F 5’-AGAGTTTGATCCTGGCTCAG-3’ and 1492R 5’-GGTTACCTTGTTACGACTT-3’ primers. The PCR reaction was as follows: 95°C for 10 min; 35 cycles of 95°C for 30 s, 55°C for 1 min, and 72°C for 1.5 min; and final extension at 72°C for 10 min. The PCR amplicon was purified to remove contaminants. Forward and reverse DNA sequencing reaction of PCR amplicon was carried out with forward primer and reverse primers using BDT v3.1 Cycle sequencing kit on ABI 3730xl Genetic Analyzer. Consensus sequence of 16S rRNA gene was generated from forward and reverse sequence data using aligner software. The 16S rRNA gene sequence was used to carry out BLAST with database of NCBI GenBank. Based on maximum identity score first ten sequences were selected and aligned using multiple alignment software program Clustal W. Distance matrix was generated and the phylogenetic tree was constructed using MEGA 7. The evolutionary history was inferred by using the Maximum Likelihood method based on the Kimura 2-parameter model.

### 2.7 Raising of antisera against purified Peptidase M84

Antiserum against Peptidase M84 was raised by immunizing a New Zealand White rabbit (bred and maintained in an animal colony as per the principles and guidelines of the ethical committee for animal care of National Institute of Cholera and Enteric Diseases))as described previously by Mondal *et al*, 2016 [41]. Intramuscular injection was given with 100 µg of purified Peptidase M84 emulsified with an equal volume of Freund’s complete adjuvant (Sigma, USA; Cat# F5881). This was followed by four booster injections with 100.0 µg of Peptidase M84 and incomplete adjuvant (Sigma, USA; Cat# F5506) at 7-day intervals. Blood samples were collected from rabbits on day 0 and 3^rd^ day after the final injection and were allowed to clot O/N at RT. Sera were collected and centrifuged (1,000 rpm, 5 min), diluted in autoclaved glycerol (Sigma, USA) and stored at −80°C until use at a dilution of 1:800 unless otherwise mentioned.

### 2.8 PCR amplification to detect the gene encodes Peptidase M84 from *Bacillus altitudinis*

Genomic DNA was isolated from 1 ml of O/N bacterial culture using Promega Wizard genomic DNA purification kit (Cat# A1120) according to the manufacturer’s protocol. 20 ng of genomic DNA was subjected to PCR amplification using GoTaq green master mix (Promega; Cat# M7122) in an automated thermal cycler (Bio-Rad, USA) to detect the presence of gene codes for Peptidase M84 using specific primers (mentioned in Table 1) under the following conditions: 10 min initial denaturation at 95 ͦ C followed by 35 cycles of 1 min denaturation at 95°C, 30 s, annealing at 55°C, followed by 1 min extension at 72°C and final 10 min extension at 72°C.

**Table 1:**
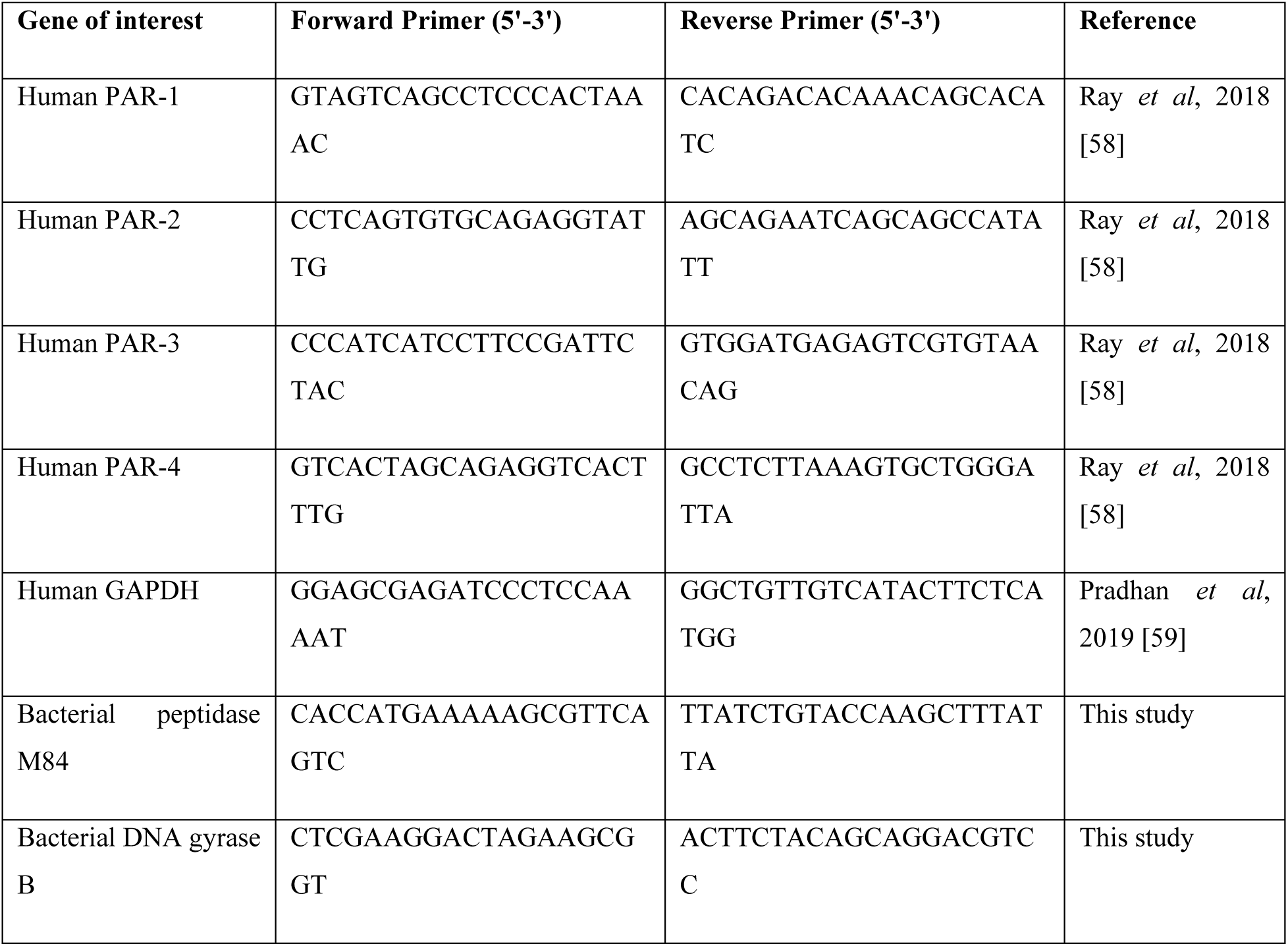
List of primers used in this study.

### 2.9 Cell culture and treatments

Human ovarian cancer cells PA-1 (ATCC; Manassas, Virginia, United State; Cat# CRL-1572, RRID:CVCL_0479) and SKOV3 (ATCC; Manassas, Virginia, United State; Cat# HTB-77, RRID:CVCL_0532) were cultured (10-12 passage) and maintained in alpha-minimum essential medium (MEM-α) (Sigma; Cat# M0644) and RPMI-1640 (Gibco; Cat# 23400021) respectively with 10% FBS, 100 U/ml penicillin G and 100 µg/ml Streptomycin sulphate at 37°C in presence of 5% CO2 incubator. Human immortalised ovarian surface epithelial cells, IOSE (IOSE-364 from N. Aueresperg and C. Salamanca, Vancouver, Canada, RRID: CVCL_5540) was maintained in MCDB-105 (Sigma Aldrich; Cat# M6395) and Medium-199 (Gibco; Cat# 31100035) in 1: 1 ratio and supplemented as stated earlier. Here, the low-passage cultures of human ovarian surface epithelium cells (isolated by scraping from human ovarian surface tissue) were immortalised by transfecting with SV40 large-T antigen viral particles [42, 43]. Mouse ovarian carcinoma ID8 (MERCK Cat# SCC145, RRID: CVCL_IU14) cell line was cultured in DMEM high glucose medium (Gibco; Cat# 31600034) supplemented with 10% FBS, 100 U/ml penicillin G and 100 µg/ml Streptomycin sulphate and 1% AOF-ITS supplement (Merck; Cat# SCM 054) at 37°C in presence of 5% CO2 incubator. All cell lines were supplied by Dr. Sib Sankar Roy (CSIR-IICB, Kolkata). Cell lines were tested for mycoplasma contamination and validated by short tandem repeat (STR) polymorphism analysis performed by the Life code genomic technologies.

### 2.10 Cell viability and cell proliferation assay

The effect of Peptidase M84 on cellular viability and to determine the sub-lethal concentration of peptidase M84, MTT assay was done as per standard protocol (Kar *et al*, 2014) [44]. About 1X10^5^ live cells/well were seeded in a 96-well plate for viability assay. Trypan blue was used to determine cell viability. After 24 h of incubation, PA-1, SKOV3 and ID8 cells were treated with different concentrations of Peptidase M84 to determine the minimal inhibitory concentration (IC50) value. After 24 h of protease treatment at different concentrations of 0.5 µg/ml – 5.0 µg/ml, MTT solution (0.8 mg/ml dissolved in serum free medium) was added to the cells and further incubated for 4 h in the dark at 37°C. The media containing MTT was removed and DMSO was added followed by incubation for 15 min in the dark. The absorbance was measured at 595 nm and mean of five replicates was taken to obtain IC50 value for subsequent experiments.

The effect of Peptidase M84 on cell viability was also studied by immunofluorescence of ki-67 nuclear antigen in PA-1 and SKOV3 cells. Peptidase M84 treated (with 2.0 µg/ml for 18 h) and untreated PA-1 and SKOV3 cells were fixed with 4 % paraformaldehyde for 10 min at RT. Cells were then permeabilised with 0.1 % Triton X-100 in 0.1% sodium citrate solution and blocked with 5% serum. Cells were incubated O/N with anti-ki-67 primary antibody (1:200) at 4°C in a moist chamber. Cells were washed with (phosphate buffer saline) PBS and then incubated with Alexa 488 conjugated secondary antibody. Nuclei were stained with DAPI for 10 min at 37°C. Cells were then washed twice with PBS, mounting was done with 10% glycerol. Images were captured using Zeiss (LSM 710) confocal microscope.

### 2.11 Flow cytometry analysis to study apoptosis

Flow cytometry was performed for screening of apoptotic activity of protease positive bacterial culture supernatants. Cellular apoptosis was detected by double staining, FITC conjugated annexinV/propidium iodide (PI) staining, as described in our previous studies (Ray *et al*, 2016) [27]. Briefly, PA-1 cells (1×10^6^) were seeded into 6 well tissue culture plates. Cells with 70% confluency were washed with PBS and starved in serum free media for 18 h. Cells were treated with filter sterilised protease positive bacterial supernatant for 18 h in complete medium. Untreated control cells were replaced with fresh media and incubated under similar conditions. After treatment, cells were harvested and flow cytometric analysis was done by using Annexin V and PI staining kit (BD, USA, Cat# 556547) as per the manufacturer’s protocol. For protease inhibition studies bacterial culture supernatant was pre-incubated with both 10 mM EDTA and 10 mM PMSF before treatment.

Flow cytometry was also performed with the purified protease as described below. After treatment with purified Peptidase M84 at a concentration range between 1.0 µg/ml to 3.0 µg/ml for 18 h, PA-1, SKOV3, IOSE and ID8 cells were harvested by centrifugation and washed twice with PBS. Cells were re-suspended in 1X binding buffer (provided with BD annexin-V kit), stained with annexin V/PI and kept at RT for 15 mins. Cells were analysed by BD FACS Aria II using ‘Cell Quest’ software. For protease inhibition studies, Peptidase M84 was preincubated with either 10 mM EDTA or 10 mM PMSF before treatment.

### 2.12 Chromatin condensation assay

After treatment with 2.0 µg/ml of protease for 18 h, ovarian cancer cells were stained with Hoechst 33342 stain (2 µg/ml) and incubated for 10 min at 37 °C, and images were taken under Zeiss confocal microscope. Condensed nucleus was counted against total number of nuclei in the field, and the percentage of apoptotic nuclei were calculated and plotted graphically [79].

### 2.13 Isolation of peritoneal exudate macrophages (PEMФ) and treatment

To assess the cytotoxic effects of purified protease on normal healthy cells of mice the peritoneal exudate macrophages (PEMФ) were isolated from 6-8 weeks of old C57BL/6 female mice as described previously (Chakraborty and Bhaumik, 2020; Naskar *et al*, 2014) [45, 46]. Briefly, Naive C57BL/6 mice were injected intraperitoneally once with 3.0 ml of a 4% (w/v) starch (Sigma, USA) solution. After 48 h, PEMФ were collected by peritoneal washing with chilled PBS followed by centrifugation of the exudate and resuspension of cell pellet in RPMI-1640 medium supplemented with 10% FBS (Gibco, USA), 100 U penicillin/ml, and 100 µg streptomycin/ml (Gibco, USA). Cells were then seeded into 6-well plates at 5×10^4^ cells/ml. The cells were then cultured for 48 h at 37°C in a humidified 5% CO2 incubator to dampen any residual effects of the starch. Non-adherent cells were removed thereafter by gentle washing with serum-free medium. The remaining adherent cells were treated with 3.0 µg/ml of Peptidase M84 for 18 h. The untreated and treated cells were collected by gentle scraping for use in the FITC conjugated annexin V/ (PI) dual staining-based apoptosis detection assay by flow cytometry as per the protocol described earlier.

### 2.14 Detection of ROS by DCFDA staining

*In-situ* ROS level was measured by oxidation of 2’,7’-dichlorofluorescin diacetate (DCFDA) to highly fluorescent 2’,7’-dichlorofluorescin (DCF). Peptidase M84 treated (2.0 µg/ml) PA-1, SKOV3, IOSE cells and ID8 (3.0 µg/ml) cells for 6 h and 18 h with their respective untreated controls were incubated with DCFDA at a working concentration of 20 µM for 20 min at 37°C, washed with PBS and subsequently the cell pellet was resuspended in 500 µl of PBS and then subjected to flow cytometry and analysed by BD FACS Aria II using ‘Cell Quest’ software. At least three independent experiments were conducted to validate our results and the mean fluorescence intensity value of DCF was plotted for quantification.

### 2.15 Flow Cytometry Detection of JC-1 Fluorescence

JC-1 dye (Invitrogen, USA) staining to detect changes in mitochondrial membrane potential in ovarian cancer cells was done according to the protocol described by Prasad *et al*, 2021 [47]. Briefly, PA-1 and SKOV3 cells (1X10^6^) were harvested by centrifugation (5 min at 500×g) after Peptidase M84 treatment (2.0 µg/ml for 6 h and 18 h). Cells were then resuspended in 500 µl of PBS. JC-1 (5,50,6,60-tetrachloro-1,10,3,30-tetraethyl-imidacarbocyanine iodide) was added to a final concentration of 10.0 µg/ml from a stock solution of 5.0 mg/ml and cells were incubated in dark at 37°C for 30 min. Cells were then washed once and again resuspended in 500 µl of PBS. Cells were analysed in a BD FACS Aria II flow cytometer (BD Bioscience, San Jose, CA, USA). The ratio of the median value of green and red fluorescence was plotted for quantification.

### 2.16 Comet assay

DNA damage (single strand breaks) was measured by alkaline comet assay (Singh *et al*., 1988) [48]. Briefly, PA-1 and SKOV3 control and Peptidase M84 treated (2.0 µg/ml for 18 h) cells were suspended in 0.6% (w/v) low melting agarose. Subsequently, cells were layered over a frosted microscopic slide previously coated with a layer of 0.75% normal melting agarose. The slides were then immersed in a lysis buffer of pH 10.0 and left overnight at 4°C. Slides were transferred into a horizontal electrophoresis chamber containing an alkaline solution (300 mM NaOH, 1 mM Na2EDTA; pH 13.0). Pre-soaking for 20 min was done in order to unwind DNA. Electrophoresis was then carried out for 20 min (300 mA, 20 V). Slides were washed thrice with neutralizing buffer (Tris Buffer 0.4 M, pH 7.5) followed by staining with ethidium bromide (final concentration 40.0 µg/ml). Slides were examined under Axio observer 7 Apotome.2; objective-EC Plan-Neofluar 40X / 0.75 NA fluorescence microscope and image analysis was done using comet assay software programme Komet 5.5. DNA damage was quantitated by tail moment measurement.

### 2.17 Western blotting (WB)

Western blot was done as described by Ray *et al*, 2016; Das *et al*, 2022 [27, 49]. Briefly, cultured cells were lysed with RIPA buffer and the concentration of protein samples was measured using BCA assay kit. Equal amount of protein was loaded onto SDS-PAGE for separation and then electrophoretically transferred to the PVDF membrane (Merck; Cat# IPVH00010). After transfer the membrane was blocked with 3% BSA (Sigma; Cat# A1470) in Tris-buffered-saline (TBS) and incubated overnight with primary antibody (1: 1000) against the protein of interest (as per requirement) at 4°C. The blot was washed with TBS-Twin 20 (TBST) buffer and incubated with HRP/AP-tagged secondary antibody for 2 h at RT. Proteins were either visualized by Bio-Rad gel documentation system using specific ECL substrate (Thermo scientific; Cat# 34580) for HRP tagged antibodies (Cell Signaling Technology Cat# 7076 and Cell Signaling Technology Cat# 7074) or using BCIP/NBT substrate (Bio-rad; Cat# 1706432) for AP tagged antibodies (Cell Signaling Technology Cat# 7056 and Cell Signaling Technology Cat# 7054).. Quantification of western blots was performed using GelQuant (Thermo Fisher Scientific, USA) and ImageJ (NIH, Bethesda, MD) software. At least three independent experiments were performed to confirm the findings and band intensities were normalized to loading controls.

### 2.18 Immunocytochemistry and confocal imaging

Peptidase M84 treated (2.0 µg/ml for 18 h) and untreated PA-1 and SKOV3 cells were fixed with 4% paraformaldehyde for 10 min at RT. Cells were then permeabilized with 0.1 % Triton X-100 in 0.1% sodium citrate solution and blocked with 5% FBS. Cells were incubated O/N with primary antibody (1:200) against the protein of interest (as per requirement) at 4°C in a moist chamber. Cells were washed with PBS and incubated with either Alexa 488 (Cell Signaling Technology Cat# 4408 and Cell Signaling Technology Cat# 4412) or Alexa 555 conjugated secondary antibody (Cell Signaling Technology Cat# 4413 and Cell Signaling Technology Cat# 4409) for 2 h at RT. Nuclei were stained with either DAPI or Hoechst 33342 (working concentration 1.0 µg/ml) for 10 min at 37°C. Cells were washed twice and mounting was done with 10% glycerol. Images were captured using confocal microscope (Zeiss 710); objective-Plan-Apochromat 63X / 1.40 NA. Fluorescence intensity and Pearson’s co-efficient values were measured manually by drawing lines along the entire length of each nucleus or cytosol and calculating them using Fiji (https://imagej.net/software/fiji/) and all the other required analysis was done according to the procedure described by Das *et al*, 2022 [49].

### 2.19 Evaluation of intracellular cytochrome c by western blot and immunofluorescence

Subcellular fractionation to extract mitochondria free cytosol from PA-1 and SKOV3 cell lysates was performed. using the method described by Dimauro *et al* 2012 [50]. Briefly, 5×10^6^ cells were harvested after 18 h of protease treatment (2.0 µg/ml) by trypsinization followed by the isolation of mitochondria free cytosolic fraction to assess the release of cytochrome c. Concentration of cytochrome c in cytosolic fraction was measured by western blot in protease treated and untreated cells. Cytosolic GAPDH (mitochondria free) was used as loading control.

To observe the intracellular cytochrome c distribution, co-localization-based immunofluorescence was used as described earlier (Li *et al*, 2020; Sun *et al*, 2008) [51, 52]. Briefly, PA-1 and SKOV3 cells were treated with Peptidase M84 for 18 h at a concentration of 2.0 µg/ml. Untreated and treated cells were incubated with Mito Tracker dye (Mito Tracker Green FM; Molecular Probes; Thermo Fisher Scientific) at a working concentration of 100 nm for 40 min in a 37°C incubator in dark. The slides were then fixed with 4% formaldehyde at RT for 15 min. The fixed slides were stained with anti-cytochrome c antibody (1:200) and kept O/N at 4°C in a moist chamber, followed by staining with Alexa 555 labelled secondary antibody (1:200) at RT for 2 h. Cells were stained with DAPI (1.0 µg/ml) for 10 min, washed twice and mounted with 10% glycerol in glass slides. Images were obtained using different excitation filters and merged. Co-localization was quantified by calculating Pearson’s co-efficient values.

### 2.20 mRNA expression level of different PARs upon Peptidase M84 treatment by quantitative Real-Time PCR (RT-qPCR)

RT-qPCR of PARs was done as described previously (Ray and Pal, 2016) [37]. Total cellular RNA was isolated from untreated and Peptidase M84 treated (2.0 µg/ml for 18 h) PA-1 and SKOV3 cells (about 1X 10^6^ cells for PA1 and 0.8X10^6^ cells for SKOV3) using RNA isolation kit (Zymo research; Cat# R1057). 2.0 µg of the total RNA was reverse transcribed using Revertaid first strand cDNA synthesis kit (Thermo scientific; Cat# K1622) to synthesize the cDNA first strand. Thereafter, the cDNA first strand was used in the subsequent amplification by Real-Time PCR with the primers described in Table 1. GAPDH was used as an internal control to normalize the results. Real-Time PCR was performed using SYBR green reagent (Eurogentec; Cat# UFRSMTBC101) in a total volume of 25 µl containing 10 pM of each primer (mentioned in Table 1), 12.5 µl of SYBR green reagent and 2.0 µl of cDNA. PCR reactions were carried out by using an ABI multicolour real time PCR detection system. The thermal cycling conditions used for Real Time PCR were: denaturation at 95°C for 30 s followed by 35 cycles of 1 min denaturation at 95° C, 30 s, annealing at 55°C, followed by 1 min extension at 72°C and final 10 min extension at 72°C.

### 2.21 Immunoprecipitation

For immunoprecipitation (IP), cells were lysed in immunoprecipitation buffer [50 mM Tris-HCl, pH 7.5, 150 mM NaCl, 0.1% Triton X-100, 1% IGEPAL, 1 mM PMSF and protease inhibitor cocktail (Sigma Aldrich)], and it was performed under denaturing conditions as described previously by Ray and Pal, 2016 [37]. Briefly, 2.0 µg/ml of Peptidase M84 was added to the PA-1 and SKOV3 cell medium and incubated for 30 min. The incubation media was removed after centrifugation and cells were fixed with 4% formaldehyde and the cell lysate was prepared using RIPA buffer. 500 µg of the total lysate was incubated with either PAR-1 antibody or antisera raised against Peptidase M84 for 5 h at 4°C in a rotating shaker at a speed of 10 rpm. The lysate along with antibody was allowed to bind with A/G agarose beads and incubated O/N at 4°C in a rotating shaker at a speed of 10 rpm. The lysates were centrifuged and washed twice with IP buffer to remove any non-specific bindings. Beads were boiled with SDS-protein loading dye for 10 min and subjected to electrophoresis. The proteins were transferred to a PVDF membrane and western blot was done with either anti-Peptidase M84 antibody (1:3000) or anti-PAR-1 antibody (1:1000). Immunoblots (IB) were developed using Veri Blot (Abcam Cat# ab131366) as per the manufacturer’s instruction to observe the interaction between Peptidase M84 and PAR-1.

### 2.22 Nuclear cytosolic fractionation

To detect the nuclear translocation of NFκB subunits, PA-1 and SKOV3 (5X10^6^) cells were treated with 2.0 µg/ml of Peptidase M84 for 18 h. Extraction of nuclear and cytosolic fractions of untreated and Peptidase M84 treated cells were performed using NE-PER™ Nuclear and Cytoplasmic Extraction Reagents (Thermo Scientific™, USA, Cat#78833) according to manufacturer’s protocol. The fractions were used for western blotting to determine nuclear translocation of p50 and p65. Here, α-tubulin and Histone H3 were used as cytosolic and nuclear loading controls respectively.

### 2.23 Treatments of cells and siRNA transfection

The cells were first starved for at least 18 h with an incomplete medium prior to respective treatments. Scrambled (SCR) siRNA (Cat#sc-37007, Santa Cruz Biotechnology) was used as control for knockdown studies. Lipofectamine 3000 was used as transfection reagent and transfection was performed according manufacturer’s protocol. The transfection was done at 50–60% confluency and the transfection reagents were added in Opti-MEM (Gibco, USA; Cat# 31985062). medium and after 4 h of transfection the media was changed to respective media of treatment. PAR-1 si-RNA (CGGUCUGUUAUGUGUCUAUdTdT) was transfected for at least 48 hours for efficient knock down.

### 2.24 Flowcytometric analysis for apoptosis and ROS detection using inhibitors to study the signalling pathway

Quantitative evaluation of apoptosis and ROS was performed using the flow cytometry methods as described previously by Ray and Pal, 2016 [37]. PA-1 and SKOV3 cells (1X10^6^ cells) were incubated with either 2.0 µg/ml of Peptidase M84 or pre-incubated for 1 h with 0.5 µM PAR1 inhibitor (ML161) or 3.0 µM of NFκB inhibitor (MG132), or 10.0 µM p38 inhibitor (SB203580) and then incubated with 2.0 µg /ml of Peptidase M84 for 18 h. Cells were washed with PBS and analysed to detect apoptosis by Annexin-V-FITC and PI dual staining based method. In-situ ROS was determined in both PAR-1 silenced and ML-161 treated cells by DCFDA staining using FACS Aria II (Cell Quest software) as per the protocol described earlier.

### 2.25 *In vivo* syngeneic mice model and ethics statement

The mice which were used in this study were healthy adult female C57BL/6 mice of approximately 22–25 g of weight (6-8 weeks old), bred and maintained in an animal colony as per the principles and guidelines of the ethical committee for animal care of National Institute of Cholera and Enteric Diseases (NICED). Survival kinetics and body weight changes were studied by implanting 5X10^6^ number of ID8 cells (0.2 ml) into the peritoneal cavity of C57BL/6 female mice and allowed to multiply [53, 54]. The day of tumour implantation was assigned as day ‘0’. In the present study, the animals were randomised into six groups containing ten animals (n = 10) in each group. (i) **Group 1** normal set (non-tumour-bearing; untreated control); (ii) **Group 2** only tumour-bearing set (ID8 control); (iii) **Group 3** Peptidase M84-treated ID8 bearing set; where 3.0 µg /ml Peptidase M84 (12.0 µg / kg of body weight) was injected intraperitonially on the day after inoculation of ID8 cells and injected once in a week for seven successive weeks; (iv) **Group 4** Peptidase M84 pre-treated with 10 mM EDTA; tumour bearing set where Peptidase M84 was inhibited with 10 mM EDTA at 37°C for 1 h and then injected into the peritoneum cavity of ID8 treated mouse for seven successive weeks (v) **Group 5** Only Buffer treated set (vehicle control) where 100 µl of 1X PBS buffer was injected intraperitonially once in a week for seven successive weeks. (vi) **Group 6** only Peptidase M84 treated group (protease control).

The life span of each group of mice was evaluated by measuring the percentage of survival rate in each group at 10 days interval by using a formula:

(Number of live animals in a group/number of initial animals in that group) X100

### 2.26 Collection of blood and serum samples

Blood samples from mice were collected as per the protocol reported previously (Barua *et al*, 2019) [55]. Before euthanasia, all animals were fasted for 4 h then the blood samples were collected by cardiac puncture into microcentrifuge tubes and left to clot. The serum was separated by centrifugation at 2000 X g for 15 min and stored at −20°C until analysis.

### 2.27 Measurement of cellular ROS and liver and kidney toxicity in mice

Cellular ROS level was detected by biochemical analysis of different markers in serum samples of all groups of experimental mice as described above. Different biochemical parameters like lipid peroxidation (LPO), reduced glutathione (GSH), catalase (CAT) and superoxide dismutase (SOD) were measured in the serum of different groups of mice at day 0-and 45-days intervals after inoculation of cells using standard protocols as described earlier by Ray *et al*, 2016 [27].

Liver toxicity markers such as serum aspartate transaminase (AST), alanine aminotransferase (ALT) level and kidney toxicity markers such as urea and creatinine were analysed by automated clinical chemistry analyzer (AU400, Olympus, Japan) according to the manufacturer’s protocol after 45 days of treatment.

### 2.28 Histology of liver and kidney tissue of mice

Histopathology and hematoxylin-eosin (H&E) staining of liver and kidney tissue was done as per protocol suggested by Dasgupta *et al*, 2022 [56]. For the experimental purpose, euthanasia was done as per CPCSEA guidelines. The liver and kidney were harvested from all six groups of mice mentioned earlier after 45 days of tumour implantation. The tissues were fixed in 10 % buffered formalin. The fixed tissue was paraffin embedded and serially sectioned at 4.0 µm, and stained with hematoxylin and eosin (H&E). Tissue Sections were viewed under 40X magnification of Zeiss Axiovert 40 C microscope.

### 2.29 *In vivo* evaluation of cell viability

The effects of 3.0 µg/ml of Peptidase M84 were compared against control groups, where mice were randomised in six different groups as mentioned earlier. After 45 days and 60 days of tumour inoculation, mice were sacrificed to collect total cells from the peritoneum and cells viability was checked by trypan blue exclusion method as described previously by Barua *et al*, 2020 [57].

### 2.30 Statistical analysis

All experiments were replicated at least thrice (n=3). All animal experimental groups contained either 10 or 6 animals. The experimental results were expressed as mean ± standard deviation. All statistical analysis was done by applying Student’s t-test (unpaired two-tailed) and bar graphs were plotted using Microsoft Office Excel 2021. In all panels, ns (non-significant) p>0.05, *p≤0.05, **p≤0.01, and ***p≤0.001. In each panel, error bars were calculated based on results obtained from a minimum of three independent experiments.

## 3. Results

### 3.1 The culture supernatant from isolate GDL-186 which showed similarity with *Bacillus altitudinis*, triggered apoptosis in PA-1 cells

High extracellular protease activity was recorded for the 12 isolates (GDL-184, GDL-185, GDL-186, GDL-187, GDL-188, GDL-191, GDL-201, GDL-71, GDL-213, BSF-4, BSF-80, BSF-32) screened out of 200 environmental isolates (Fig. 1A, B). Initially, we assessed the apoptosis inducing potentials of the sterile supernatants extracted from these isolates among PA-1 cells by flow cytometry after treatment. Herein, PA-1 cells, specifically treated with GDL-186 supernatant underwent apoptosis significantly (Fig. 1C, D). This was reconfirmed when the GDL-186 supernatant also successfully induced apoptosis in SKOV3 cells additionally. Contrarily, no apoptosis inductions were noted among both PA-1 and SKOV3 cells upon treatment with GDL-186 supernatant pre-incubated with EDTA and PMSF. A significant decrease in the percentage of apoptotic PA-1 and SKOV3 cells was enumerated, denoting protease mediated apoptosis inducing property of GDL-186 isolate (Supplementary Fig. S1A-D). Thereafter, we identified GDL-186 as a *Bacillus* species after deciphering 100% sequence homology of their 16S rRNA with that of *Bacillus altitudinis* strain in NCBI Blast results. The gene sequence was submitted to NCBI GenBank under accession number OP738003.1. (Supplementary Fig. S1E, F).

**Fig. 1.**
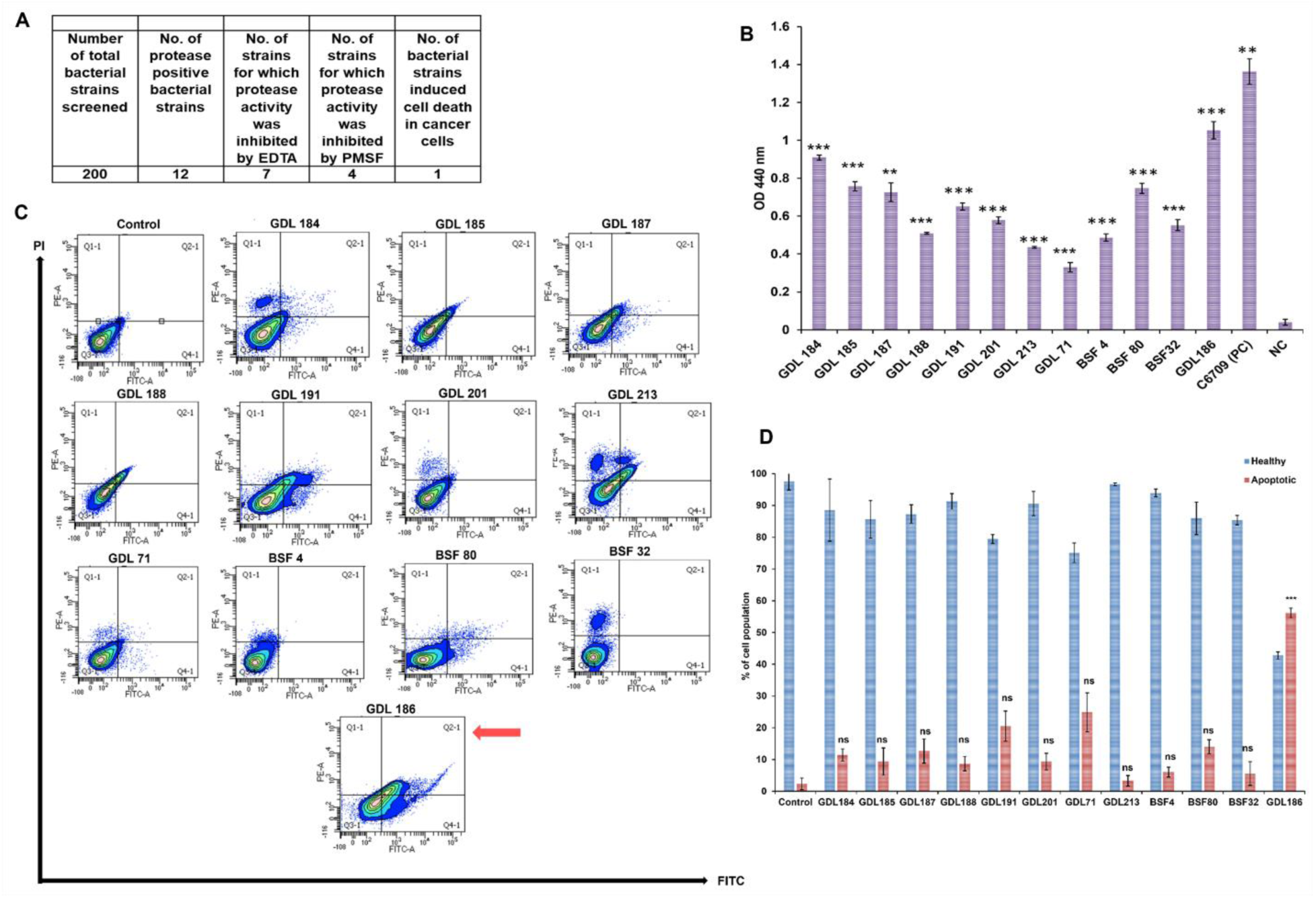
Screening of 200 bacterial isolates from environmental sources for extracellular protease activity and apoptosis inducing activity. **A, B** Results of azocasein (0.7%) assay are represented in tabular and graphical form. Culture supernatant of *Vibrio cholerae* EL Tor strain C6709 was used as a positive control and nutrient broth was used as the negative control. **C** Flow-cytometric analysis to assess apoptosis was performed with the culture supernatant of the protease-secreting isolates in PA-1 cells. In each display the lower right quadrant is for early apoptotic cells (Annexin V+/PI-), the upper right is for late apoptotic cells (Annexin V+/PI+), the upper left is for necrotic population (Annexin V-/PI+) and lower left is for healthy cells (Annexin V-/PI-). The culture supernatant of GDL-186 isolate (marked with an arrow) showed significant apoptosis. **D** The above results are graphically represented in the bar diagram. All statistical analysis was done by applying the Student’s t-test (unpaired two-tailed). Data are expressed in ± SEM. In all panels, (non-significant) ns p>0.05, *p≤0.05, **p≤0.01, and ***p≤0.001. Error bars were calculated based on results obtained from a minimum of three independent experiments (n=3).

### 3.2 Purification and identification of secreted protease named Peptidase M84 from *Bacillus altitudinis* GDL-186

GDL-186 culture supernatant proteolytically degraded azocasein alongside inducing apoptosis in PA-1 cells. On this basis, we decided to purify and characterise this supernatant for its typical protease-like behaviour. Firstly, EDTA and 1, 10-phenanthroline treatment inhibited the proteolytic activity of these crude supernatants which remained unaltered in PMSF and EGTA’s presence. This hinted at GDL-186 being an extracellular zinc-dependent metallo-protease (Supplementary Fig. S1G, H). Next, the probable protease was concentrated following dialysis where the resultant non-binding and binding fractions were examined for proteolytic activity (Fig. 2A, Supplementary Fig. S1I) Interestingly, the non-binding fraction was observed to exhibit protease activity (Fig. 2B). This was therefore pooled, concentrated and analysed by gel filtration (sephadex G-75). Two fractions (indicated as two different peaks in the chromatogram) were eluted from the G-75 column of which the first fraction showed higher protease activity compared to the second (Fig. 2C, D). The first peak of sephadex G-75 elution was hence pooled, concentrated and further analysed in SDS-PAGE (15%). Two major bands around 25 kDa and 16 kDa molecular weights were distinctly observed which upon EDTA pre-incubation resulted in a single band at 25 kDa only (Fig. 2E). This fraction of G-75 also showed single band in Native-PAGE (10%) (Fig. 2F). These bands were further characterised by nano-LC-MS/MS-TOF where the generated peptide sequence was homologous to ‘**Peptidase M84**’ from *Bacillus altitudinis*. (Fig. 2G). This enzyme typically consists of a consensus amino acid sequence HExxH and a Met-turn motif ‘CLMNY’ in downstream of its active site. The histidines and glutamic acid act as zinc ligands and catalytic base respectively [60].

**Fig. 2.**
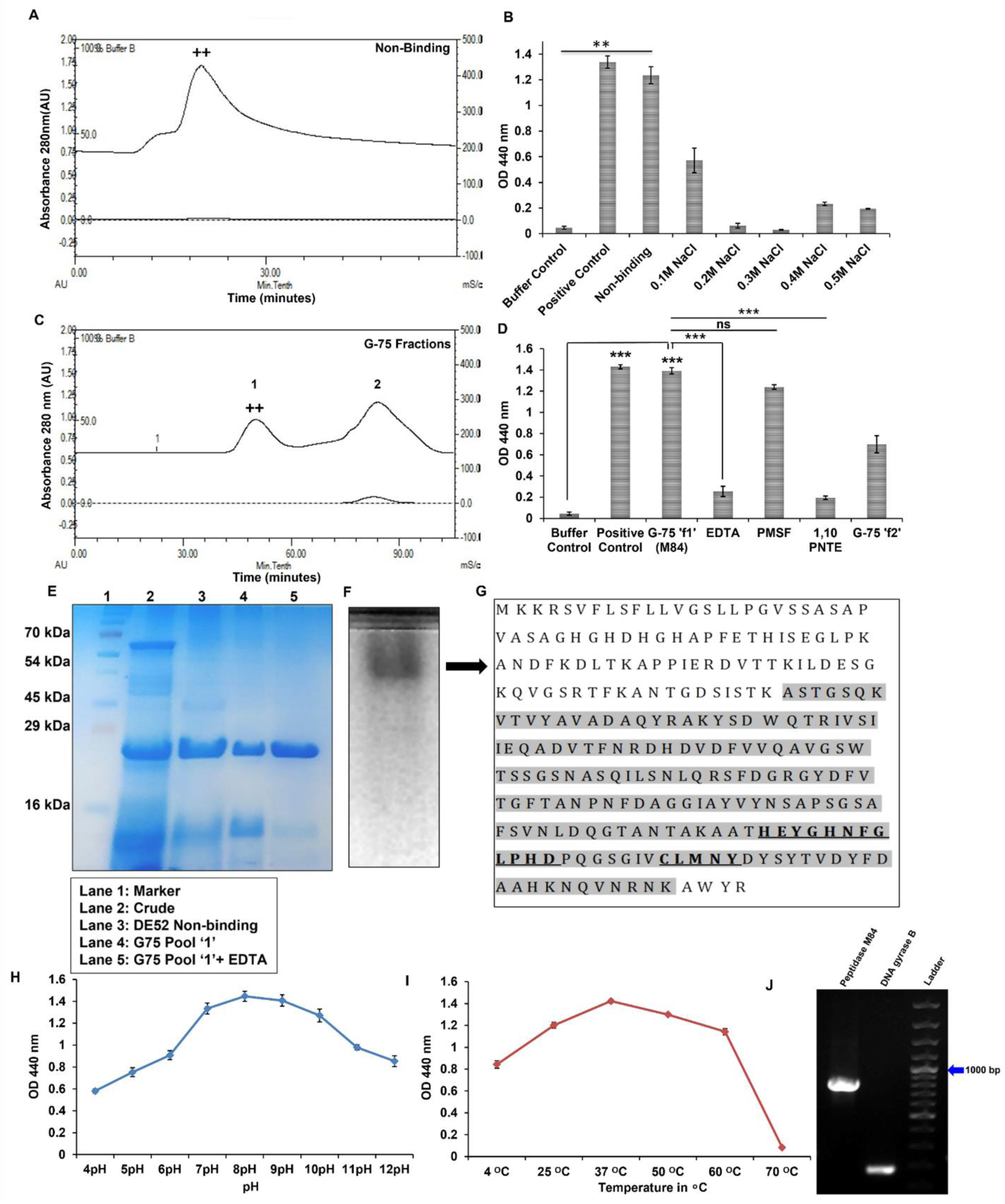
Purification and identification of Peptidase M84 from *Bacillus altitudinis* GDL-186. **A** Chromatogram of anion exchange chromatography of the ammonium sulphate precipitated crude protein fraction from GDL-186 culture supernatant. **B** Azocasein assay performed with protein fractions (5.0 μg) obtained from DEAE-52 column showed higher protease activity in the non-binding fraction (++) as compared to the NaCl (0.1 M - 0.5 M) eluted binding fractions. **C** Chromatogram of the G-75 gel filtration chromatography showed separation of the non-binding fraction, represented as two individual peaks (G-75’f1’ and G-75’f2’). **D** Azocasein assay with G-75 fractions showed maximum protease activity in G-75 ‘f1’ (++) fraction. This activity was inhibited by EDTA and 1,10-phenanthroline. **E** 15% SDS-PAGE profile of the purified protease showed two major bands near 25 kDa and 16 kDa, which upon pre-incubation with EDTA showed a single band at 25 kDa. **F** 12% Native-PAGE profile of the purified protease (G-75 ‘f1’) showed a single band. **G** Bands of the protease from both native and SDS-PAGE were identified by Nano-LC-MS/MS-TOF peptide sequencing which exhibited homology with ‘Peptidase M84’. The peptides showing homology are with the background colour. The active sites and the zinc-binding domains of the protease are underlined. **H, I** Determination of the optimum pH and temperature of Peptidase M84. **J** PCR amplification of the full-length (813bp) peptidase M84 gene (Lane 1) and DNA gyrase B gene (Lane 2) from *Bacillus altitudinis* strain GDL-186. All statistical analysis was done by applying Student’s t-test (two-tailed). Data are expressed in ± SEM. In all panels, ns p>0.05, *p≤0.05, **p≤0.01, and ***p≤0.001. In each panel, error bars were calculated based on results obtained from a minimum of three independent experiments (n=3).

### 3.3 Inhibition of the protease activity revealed the metallo-protease nature of Peptidase M84 which was optimally active at normal physiological pH and temperature

Purified Peptidase M84 was also proteolytically active against azocasein. This activity was yet again inhibited by EDTA and 1,10-phenanthroline but remained unaltered with PMSF (Fig. 2D). Inhibition studies once again confirmed that this purified Peptidase M84 was a zinc-containing metallo-protease. Additionally, native gelatin zymogram profile revealed Peptidase M84 to be proteolytically active against gelatin. A clear hollow zone due to gelatin degradation was attained in Native-PAGE (with 0.1% gelatin) followed by coomassie staining (Supplementary Fig. S1J). Peptidase M84 showed proteolytic activity over a wide range of pH values (from 4.0–11.0). Although, the optimal activity and the minimal activity of Peptidase M84 were recorded at pH 8.0 and pH 4.0 respectively. The enzyme kept 80–90% activity at pH 7.0–10.0 (Fig. 2H). In line with this, Peptidase M84 exhibited maximum activity against azocasein at temperatures ranging from 37°C to 40°C which decreased gradually at 50°C to 60°C. This protease showed suboptimal activity at temperatures ranging from 4°C to 25°C. At higher temperature ranges of 60°C to 70°C, the proteolytic activity was almost completely abolished (Fig. 2I). Besides, the full-length amplified 813 bp gene encoded Peptidase M84 of *Bacillus altitudinis* was also detected in 0.8 % agarose gel after PCR amplification (Fig. 2J)

### 3.4 Peptidase M84 exhibited apoptosis and suppressed the proliferation of ovarian cancer cells but had no such impact on IOSE and PEMФ cells

MTT cell viability assay showed a gradual depletion in the percentage of viable ovarian cancer cells with increasing concentrations of Peptidase M84, ranging between 0.5 µg/ml – 5.0 µg/ml. The *in vitro* safe dose (IC50) of Peptidase M84 was found to be 2.0 µg/ml for PA-1 and SKOV3 cells. Furthermore, Peptidase M84 showed IC50 3.0 µg/ml against ID8 cells (Fig. 3A-C). These concentrations were in line with the concentrations used to detect apoptosis and to study the associated bio signalling pathways in the aforementioned cell lines.

**Fig. 3.**
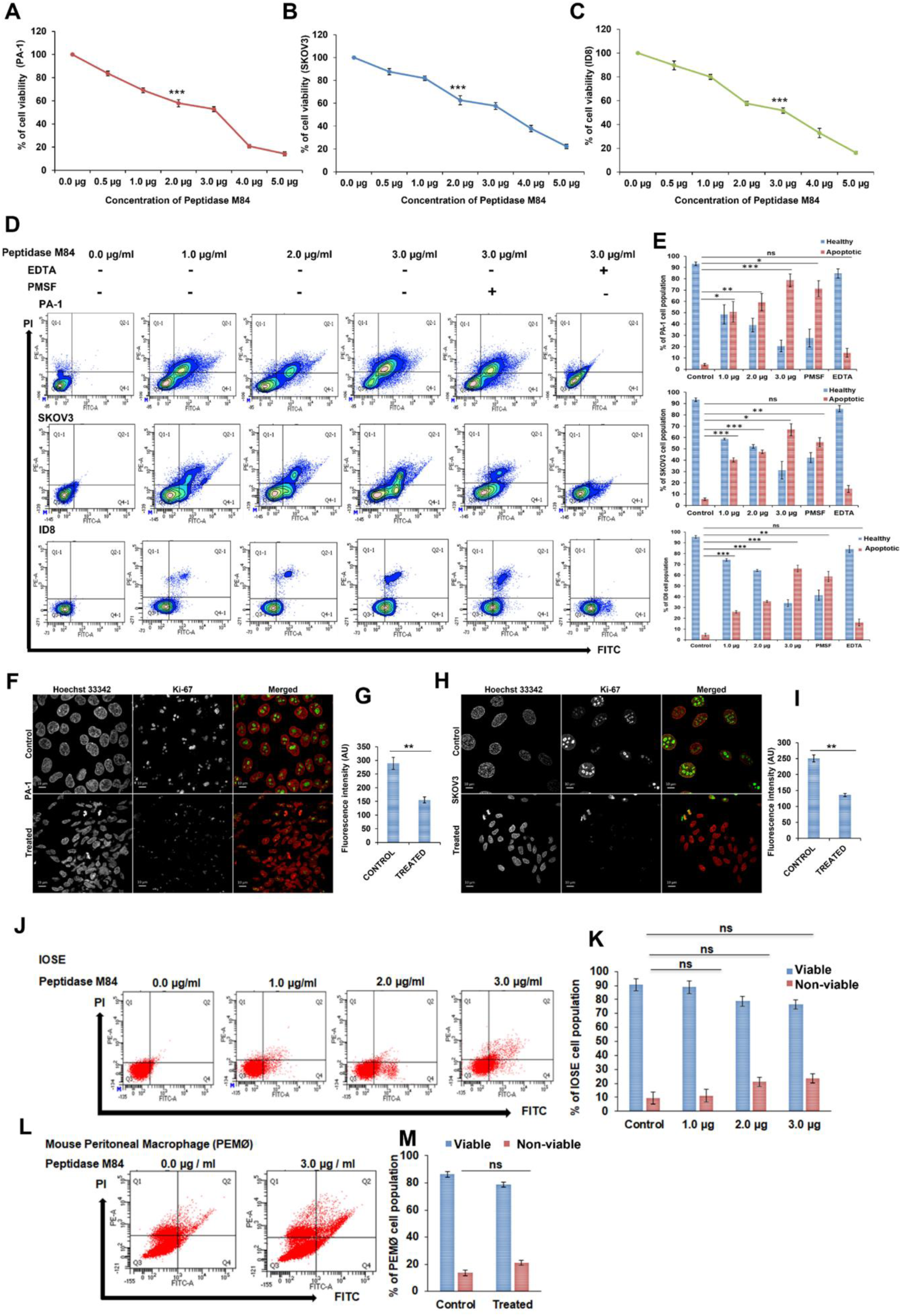
Detection of cytotoxicity and apoptosis in Peptidase M84 treated ovarian cancer cells and human ovarian normal epithelial cells (IOSE) and in mouse peritoneal macrophage (PEMФ) cells. **A-C** MTT assay revealed the IC50 value of Peptidase M84 in PA1, SKOV3 and ID8 cells. **D** Flow cytometric analysis of Peptidase M84 treated PA1, SKOV3 and ID8 cells showed dose dependent apoptosis. EDTA inhibition significantly reduced the effect while PMSF treatment did not show any significant effect on cell viability. **E** Bar diagrams show the percentage of healthy and apoptotic cell population. **F-I** Confocal imaging of nuclear Ki-67 (green) in PA1 and SKOV3 cells along with the bar diagrams of mean fluorescence intensity showed a significant reduction in cancer cell proliferation due to Peptidase M84 treatment. **J-M** Flow cytometric analysis of the purified Peptidase M84 treated IOSE and PEMФ cells showed no significant apoptosis as compared to the untreated cells in similar doses. Bar diagrams of the data are also being presented. All statistical analysis was done by applying Student’s t-test (two-tailed). Data are expressed in ± SEM. In all panels, ns p>0.05, *p≤0.05, **p≤0.01, and ***p≤0.001. In each panel, error bars were calculated based on results obtained from a minimum of three independent experiments (n=3). Scale bars: 10 µm.

Next, in order to investigate the cytotoxic effect of Peptidase M84, PA-1, SKOV3 and ID8 cells were treated with doses ranging between 1.0 μg/ml to 3.0 μg/ml for 18 h. Flow cytometric analysis showed that Peptidase M84 caused significant apoptosis in these cell lines as compared to untreated cells. Inhibition of protease activity by EDTA showed a significant reduction in the percentage of apoptotic cells. However, this percentage was not affected by PMSF. These results clearly indicated that purified Peptidase M84 by virtue of its metalloprotease activity could promote cell death by inducing apoptosis in human and mouse ovarian cancer cells (Fig. 3D-E).

Ki-67 protein is considered as a proliferation marker for human tumour cells. Higher expression of nuclear Ki-67 is found in all stages of cell cycle not including the G0 stage and dead cells [61]. We noticed a lower abundance of Ki-67 in Peptidase M84 treated PA-1 and SKOV3 cells as compared to the untreated control cells indicating that Peptidase M84 significantly restrained cancer cell proliferation. Contrarily, higher levels of nuclear Ki-67 in untreated control cells indicated rapid proliferation and increased survivability (Fig. 3F-I).

Peptidase M84 treatment also altered the morphology of PA-1 and SKOV3 cells. Peptidase M84 showed a cell distending effect at lower concentrations. A cell rounding effect was observed when cells were treated with increasing concentrations of Peptidase M84 for 18 h (Supplementary Fig. S2A).

During apoptosis, highly condensed inert and fragmented chromatin gets packaged into apoptotic bodies [41]. Herein, these became distinct upon Hoechst 33342 staining which indicated apoptotic induction. In Peptidase M84 treated PA-1, SKOV3 and ID8 cells, the percentage of condensed nuclei were 43.66, 38.33 and 44.33% respectively whereas the respective control cells were shown to be 13%, 10.6%, and 11.33% (Supplementary Fig. S2B, C). The result indicated that nuclei of Peptidase M84 treated cells exhibited condensed and bright nuclei while the untreated control cells showed less amount of condensed chromatin.

To understand the specificity, the effect of Peptidase M84 was evaluated on IOSE cells and PEMФ cells. Flow cytometric analysis showed no significant apoptosis in IOSE cells and PEMФ cells in a similar range of doses of Peptidase M84. Almost all cells remained viable even at a concentration of 3.0 μg/ml of Peptidase M84. These findings indicated that purified Peptidase M84 selectively triggered apoptosis in malignant ovarian cells but not in normal cells (Fig. 3J-M)

### 3.5 Peptidase M84 augmented ROS generation and activated the intrinsic canonical pathway of apoptosis in ovarian cancer cells

Excessive production of reactive oxygen species overtakes the cellular antioxidant capacity. It results in a state of oxidative stress, which ultimately contributes to the regulation of a wide range of signalling pathways and can impart oxidative damage mediated cell death [62, 63]. The oxidative stress related to Peptidase M84 treatment was monitored by staining with DCFDA. Under the stimulation of ROS, DCFH is converted to fluorescent DCF upon oxidation. Peptidase M84 treatment showed an increase in ROS generation in PA-1, SKOV3 and ID8 cells, depicted as an increase in the mean fluorescence intensity level of DCF in comparison to untreated cells with time (6 h and 18 h). This could lead to initiating apoptotic signals (Fig. 4A, B). In contrast, Peptidase M84 failed to promote ROS generation in IOSE cells (Supplementary Fig. S2D, E).

**Fig. 4.**
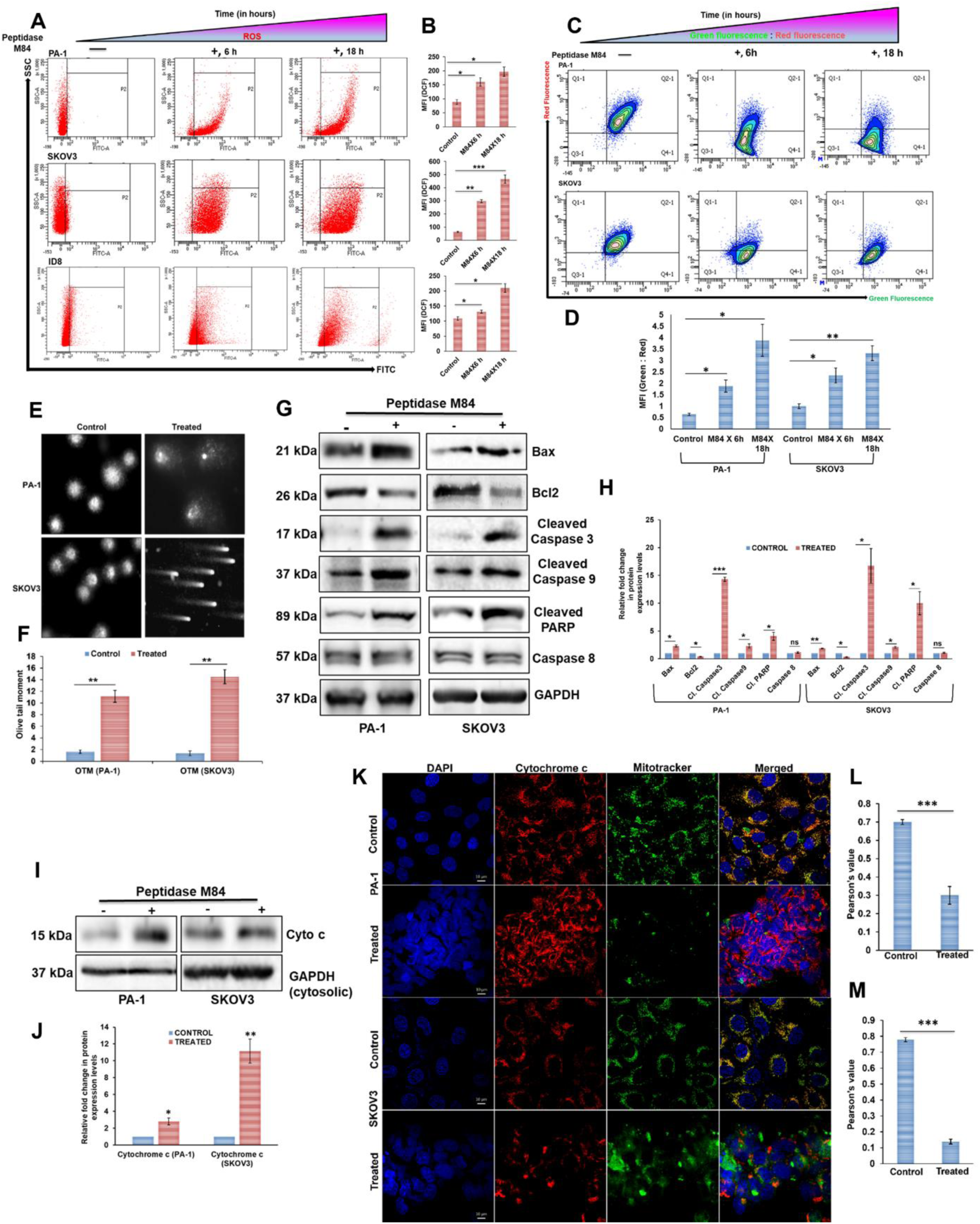
Ovarian cancer cells undergo ROS dependent mitochondria mediated intrinsic pathway due to Peptidase M84 treatment. **A** An increase in ROS generation in PA-1, SKOV3 and ID8 cells after Peptidase M84 treatment was observed by flow cytometry in a time dependant manner (6 h and 18 h). Quadrant ‘P2’ represents cells producing ROS. **B** Bar diagrams represent the mean fluorescence intensity of ROS generation. **C, D** JC-1 staining data with bar diagrams of mean fluorescence intensity (green: red) showed the disruption of mitochondrial membrane potential with time (6 h and 18 h) in Peptidase M84 treated PA-1 and SKOV3 cells. An increase in the cell population positive for green fluorescence signified disrupted mitochondrial membrane potential. **E** Alkaline comet assay represented an increase in the comet tail length in Peptidase M84 treated PA-1 and SKOV3 cells. **f** Bar diagram represents the olive tail moment of cells. **G, H** Western blot and densitometric analysis revealed alterations in expression levels of major proteins associated with the apoptotic pathway in Peptidase M84 induced and un-induced PA-1 and SKOV3 cells. **I, J** Western blot analysis with densitometry scan of cytochrome c in the mitochondria free cytosolic fraction in Peptidase M84 treated PA-1 and SKOV3 cells depicted the release of cytochrome c from mitochondria. **K-M** Immunofluorescence imaging along with bar diagrams of Pearson’s co-efficient values showed reduction in co-localization between mitochondria (green) and cytochrome c (red) and also displayed the release of cytochrome c from damaged mitochondria to cytosol due to Peptidase M84 treatment in PA-1 and SKOV3 cells. In each panel, error bars were calculated based on results obtained from a minimum of three independent experiments. All statistical analysis was done by applying Student’s t-test (two-tailed). Data are expressed in ± SEM. In all panels, ns p>0.05, *p≤0.05, **p≤0.01, and ***p≤0.001. Scale bars: 10 µm.

JC-1 dye uptake pattern suggested that Peptidase M84 also caused alterations in mitochondrial membrane potential (ΔΨm) in PA-1 and SKOV3 cells. A decrease in ΔΨm is considered as a major hallmark of apoptosis [64]. An increase in the ratio of green fluorescence to red fluorescence intensity levels clearly indicated disrupted ΔΨm in Peptidase M84 treated cells in a time dependent manner (6 h and 18 h) as compared to the untreated control cells (Fig: 4C, D). Herein, comet assay was performed to detect DNA damage in PA-1 and SKOV3 cells as a result of Peptidase M84 treatment as genotoxicity tests predominantly contribute to cancer research. We observed that Peptidase M84 treatment for 18 h gave a prominent induction of comet tail moment in ovarian cancer cells with respect to control untreated cells (Fig. 4E, F).

Next, we examined the quantitative expression profiles of some prime regulatory proteins to study the possible molecular mechanism of apoptotic cascade induced by Peptidase M84. We observed that Peptidase M84 treatment resulted in the down-regulation of anti-apoptotic Bcl-2 levels and upregulation of pro-apoptotic marker Bax in PA-1 and SKOV3 cells concerning the untreated cells. Our results also revealed activation of caspase 9, caspase 3 and upregulation of cleaved PARP expression levels by Peptidase M84. However, a consistent profile of caspase 8 expression levels indicated non-involvement of extrinsic (FADD-Caspase 8) apoptosis pathway in response to Peptidase M84 treatment in these cells (Fig. 4G, H). Interestingly, no activation of caspase 3 was observed in Peptidase M84 treated normal IOSE cells (Supplementary Fig. S2F, G)

Mitochondrial cytochrome c functions as an electron carrier in the respiratory chain [65], translocates to the cytosol in Peptidase M84 treated cells undergoing apoptosis, where it participates in the activation of specific caspases. We found an increase in the cytochrome c expression levels in cytosolic fraction devoid of mitochondria in translational levels in Peptidase M84 treated PA-1 and SKOV3 cells compared to the untreated cells. Densitometric data showed that the increase was highly significant. (Fig. 4I, J).

Immunofluorescence data also supported this observation. Damaged mitochondria due to Peptidase M84 treatment were observed with mitotracker green FM staining. We also noticed a depletion in green fluorescence intensity levels in these cells. There was a reduction in co-localisation between cytochrome c and mitochondria in PA-1 and SKOV3 cells treated with Peptidase M84 as compared to the untreated cells. Released cytochrome c was found to be distributed in punctate (red) form throughout the cytosol and around the nucleus of cells. Pearson’s co-efficient values showed a significant reduction in co-localisation between mitochondria and cytochrome c in Peptidase M84 treated PA-1 and SKOV3 cells (Fig: 4K-M).

Taken together, our findings implicated the activation of the intrinsic pathway of apoptosis in ovarian cancer cells by Peptidase M84 from *B.altitudinis*.

### 3.6 Peptidase M84 specifically interacted with PAR-1 and caused overexpression of PAR-1 in human and mouse ovarian cancer cells however not in human healthy ovarian epithelial cells

Real-Time PCR data represented overexpression of PAR-1 in PA-1 and SKOV3 cells. We found almost 4-fold and 7-fold upregulation of PAR-1 mRNA levels in PA-1 and SKOV3 cells respectively as compared to the other PARs. (Fig. 5A).

**Fig. 5.**
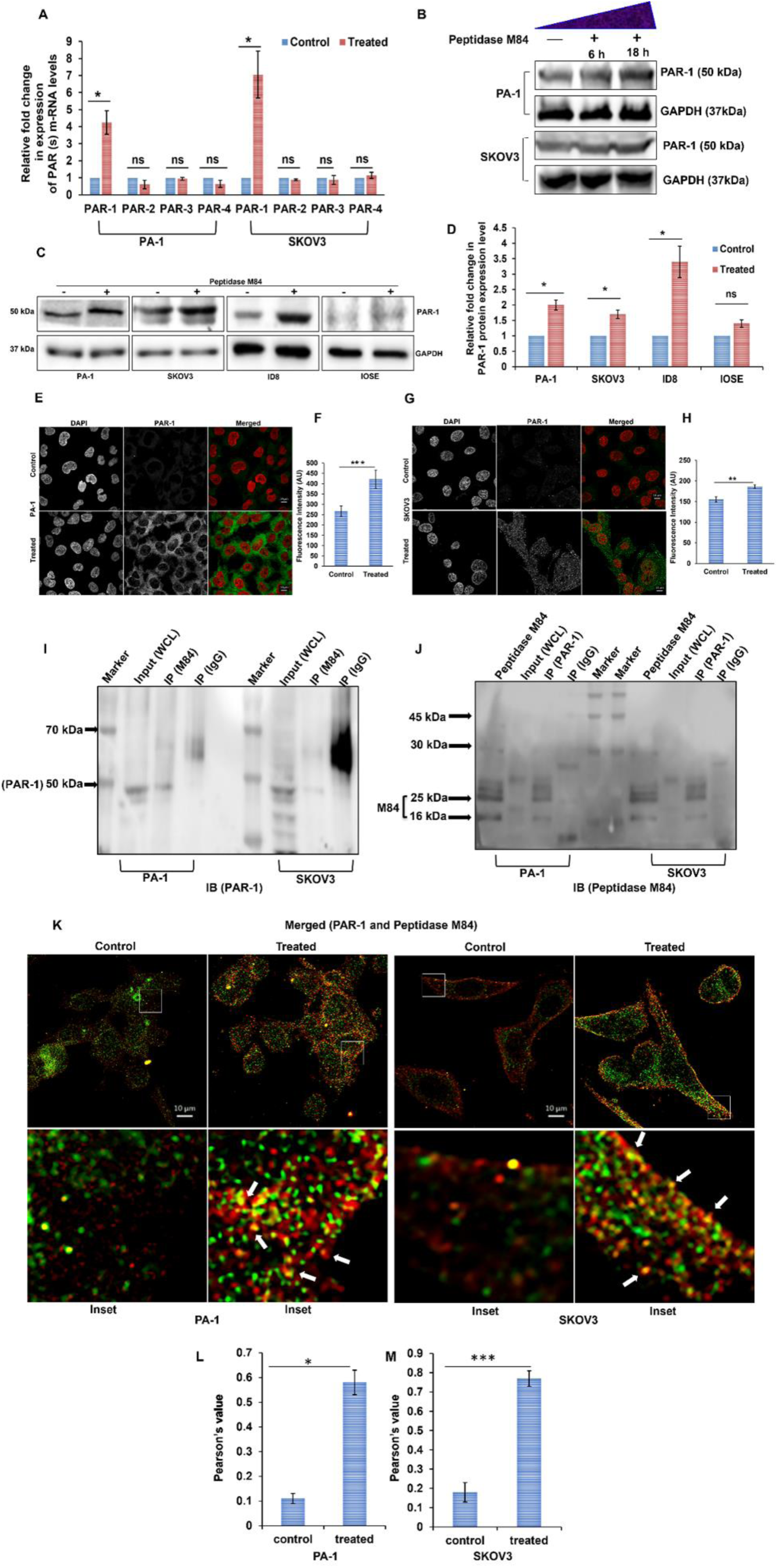
Peptidase M84 persuaded apoptotic activity in ovarian cancer cells by modulating PAR-1. **A** Real time PCR analysis of PAR-1, PAR-2, PAR3, and PAR-4 genes in Peptidase M84 treated and untreated PA1 and SKOV3 cells. Ct values are normalized to GAPDH expression, and 2^-ddct^ values were calculated (n = 3). **B** Western blot showed a time dependent (6 h and 18 h) increase in PAR-1 expression levels in Peptidase M84 treated PA-1 and SKOV3 cells. **C, D** Western blot and bar diagram of densitometry scan of PAR-1 expression in Peptidase M84 treated and untreated (18 h) PA-1, SKOV3, ID8 and IOSE cells. **E-H** Confocal imaging with respective bar diagrams of mean fluorescence intensity of PAR-1 protein expression levels (green) in PA-1 and SKOV3 cells. **I** Co-immunoprecipitation study of PAR-1 and Peptidase M84 revealed the interaction between PAR-1 and Peptidase M84 in both PA-1 and SKOV3 cells. Here, whole cell lysates (WCL) from these cells were loaded as inputs. **J** Bands of Peptidase M84 were observed in IB when Peptidase M84 was immunoprecipitated with anti-PAR-1 antibody. Here, purified Peptidase M84 from *B. altitudinis* was loaded as input. **K** Confocal microscopic imaging revealed co-localisation between Peptidase M84 (green) and PAR-1(red) in Peptidase M84 treated PA-1 and SKOV3 cells as compared to untreated cells. **L, M** Bar diagrams revealed the Pearson’s co-efficient values. In each panel, error bars were calculated based on results obtained from a minimum of three independent experiments. All statistical analysis was done by applying the Student’s t-test (two-tailed). In all panels, ns p>0.05, *p≤0.05, **p≤0.01, and ***p≤0.001. Data are expressed in ± SEM. Scale bars: 10 µm.

A time dependent (6 h and 18 h) overexpression of PAR-1 was observed in PA-1 and SKOV3 cells after Peptidase M84 treatment (Fig. 5B). Immunoblots also confirmed almost 2-fold increase in PAR-1 expression levels in PA-1, SKOV3 cells and nearly 3.5-fold overexpression in ID8 cells after Peptidase M84 treatment (18 h) (Fig. 5C, D). Importantly, no significant change was observed in the PAR-1 expression in IOSE cells (Fig. 5C, D). Almost no expression of PAR-1 was observed in IOSE cells. Densitometric scan analysis depicted the intensity differences between PAR-1 expression levels in Peptidase M84 treated and untreated cells (Fig. 5D). Immunofluorescence imaging also showed a significant increase in PAR-1 expression due to Peptidase M84 treatment in PA-1 and SKOV3 cells (Fig. 5E-H). This supports the fact that Peptidase M84 unable to induce apoptosis in normal cells via effective modulation of PAR-1.

Next, we proceeded with immunoprecipitation experiments to identify the receptors of Peptidase M84 and to elucidate the physical interaction between them. The outcome revealed Peptidase M84 specifically bound and interacted with PAR-1 when PA-1 and SKOV3 cells. A band of PAR-1 (near 50 kDa) was observed in western blot when PAR-1 was immunoprecipitated with Peptidase M84 specific antisera (Fig. 5I). Additionally, bands of Peptidase M84 (25 kDa and 16 kDa) were also observed when Peptidase M84 was immunoprecipitated with anti-PAR-1 antibody (Fig. 5J). In this context, another western blot with the raised antisera against Peptidase M84 showed two major bands at 25 kDa and 16 kDa. These bands were similar to that in 15% SDS-PAGE profile of purified Peptidase M84 (Supplementary Fig. S2 H). Furthermore, binding and interaction between Peptidase M84 and PAR-1 were also validated by co-localisation based confocal imaging. An increase in Pearson’s co-efficient values in Peptidase M84 treated (30 min) PA-1 and SKOV3 cells as compared to untreated cells indicated the possible interaction between PAR-1 and Peptidase M84 (Fig. K-M).

### 3.7 Peptidase M84 induced activation of nuclear factor κB (NFκB) and modulation of MAP kinase pathways in ovarian cancer cells

Thrombin-mediated PAR-1 activation regulates major signalling pathways such as p38 MAPK and NFκB which ultimately triggers further ROS generation. Immunoblots of cytosolic and nuclear fractions of PA-1 and SKOV3 cells showed Peptidase M84 treatment promoted nuclear translocation p50 and p65 proteins, suggesting the activation of NFκB signalling. We observed significant expression of p50 and p65 proteins in the nuclear fraction of Peptidase M84 treated cells as compared to untreated cells. Densitometry represented the ratio of protein expression between nucleus to cytosol of Peptidase M84 induced and un-induced cells. Histone H3 (nuclear protein) and alpha-tubulin (cytosolic protein) were used as loading controls (Fig. 6A, B) Immunofluorescence imaging of Peptidase M84 treated and untreated cells further supported our findings and showed prominent nuclear translocation of the same proteins. There was an increase in the ratio of nuclear to cytosolic mean fluorescence intensity of p50 and p65 proteins in Peptidase M84 treated cells (Fig. 6C-J).

**Fig. 6.**
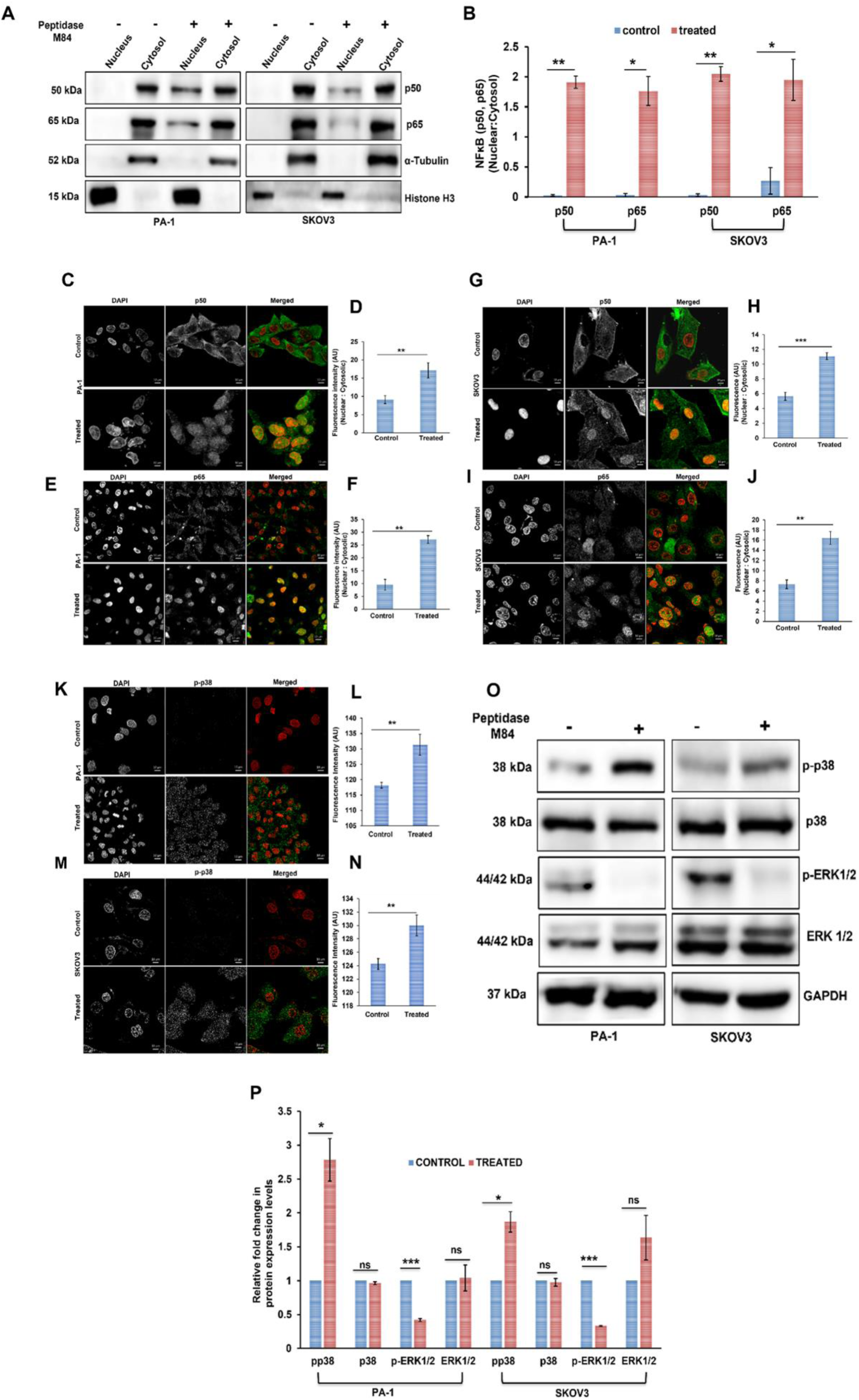
Effect of Peptidase M84 on NFκB and MAP-Kinase signalling pathways in PA1 and SKOV3 cells. **A, B** Western blot and bar diagram of densitometric scan analysis (nucleus: cytosol) indicated enhanced nuclear translocation of p50 and p65 proteins after Peptidase M84 treatment in PA-1 and SKOV3 cells for 18 h. **C-J** Immunofluorescence images showed Peptidase M84 induced nuclear translocation of p50 (green) and p65 (green) in PA-1 and SKOV3 cells. Bar diagrams represented ratio of the fluorescence intensity of nucleus to cytosol of cells. **K-N** Immunofluorescence imaging and bar diagram of mean fluorescence intensity are showed Peptidase M84 mediated increase in phospho-p38 expression level (green) in PA-1 and SKOV3 cells. **O, P** Western blot with bar diagram of densitometric scan showed significant upregulation in phospho-p38 and down-regulation phospho-ERK-1/2 expression levels in Peptidase M84 treated PA-1 and SKOV3 cells as compared to the untreated cells. In each panel, error bars were calculated based on results obtained from a minimum of three independent experiments. All statistical analysis was done by applying Student’s t-test (two tailed). In all panels, ns p>0.05, *p≤0.05, **p≤0.01, and ***p≤0.001. Data are expressed in ± SEM. Scale bars: 10 µm.

Immunofluorescence indicated the overall increase of p-p38 expression levels in Peptidase M84 treated cells (Fig. K-N). Results obtained from immunoblot also depicted the level of phospho-p38 and phospho-ERK1/2 was significantly increased and decreased respectively due to Peptidase M84 treatment. However, the entire cellular p38 and ERK1/2 levels remained unaltered in Peptidase M84 treated cells as compared to untreated cells. Thus, our report revealed the Peptidase M84 also modulated MAP kinase pathways in ovarian cancer cells (Fig. 6O, P).

### 3.8 Signalling inhibition and gene silencing studies confirmed that Peptidase M84 persuaded apoptotic signalling via modulation of PAR-1, ROS, MAP kinase and NFκB mediated axis in ovarian cancer cells

We aimed to decipher whether the interaction between Peptidase M84 and PAR-1 together with the modulation of ROS, MAP kinase and NFκB directly persuaded apoptotic signalling in ovarian cancer cells. Peptidase M84 treatment on PA-1 and SKOV3 cells showed cellular apoptosis. When these cells were priorly treated with PAR-1 inhibitor (ML161) followed by treatment with Peptidase M84 for 18 h, there was an absence of apoptosis. Furthermore, knockdown of PAR-1 was observed by si-RNA transfection for 48 h in PA-1 and SKOV3 cells (7A, B). We also found significant inhibition of apoptosis when PAR-1 silenced cells were treated with Peptidase M84 as compared to the untreated and only Peptidase M84 treated cells. PAR-1 silenced cells behaved like untreated cells even after Peptidase M84 treatment. Hence, our data illustrated that Peptidase M84 induced apoptosis in ovarian cancer cells in a PAR-1 dependent manner (Fig. 7C-E).

**Fig. 7.**
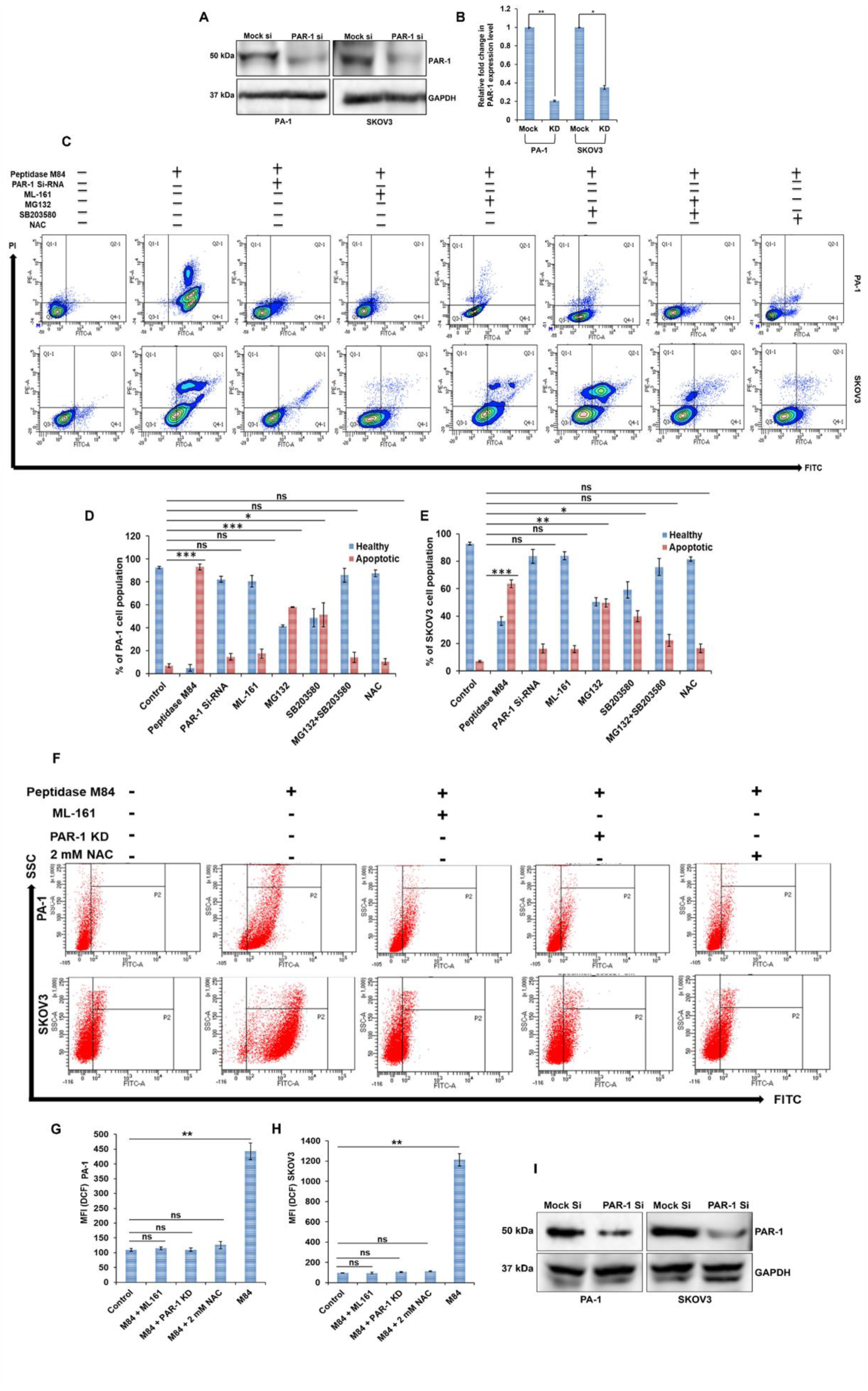
Effect of different signalling inhibitors on apoptosis mediated by Peptidase M84. **A, B** Western blot and bar diagram of densitometric scan showed knock down of PAR-1 protein expression after 48 h of si-RNA treatment in PA-1 and SKOV3 cells. **C-E** Peptidase M84 mediated cellular apoptosis was analysed by FACS in presence and absence of different signalling inhibitors (NFκB, p38, ROS and PAR-1) and PAR-1 si-RNA in PA-1 (upper panel) and SKOV3 (lower panel) cells. The above results are graphically represented in the bar diagram. **G, H** Inhibition of the PAR-1 expression by si-RNA and suppression of PAR-1 activity by ML161 resulted in inhibition of ROS generation in PA-1 and SKOV3 cells even after Peptidase M84 treatment. Cells pre-treated with 2 mM NAC were used as a negative control. Bar diagrams represented the mean fluorescence intensity of DCF. **I** knock down of PAR-1 protein expression after 48 h of si-RNA treatment in PA-1 and SKOV3 cells for the previous experiment. All statistical analysis was done by applying Student’s t-test (two-tailed). In all panels, ns p>0.05, *p≤0.05, **p≤0.01, and ***p≤0.001. Data are expressed in ± SEM. In each panel, error bars were calculated based on results obtained from a minimum of three independent experiments.

Peptidase M84 induced activation of MAP kinase and NFκB and also augmented ROS levels in ovarian cancer cells. To confirm whether these signalling pathways were directly responsible for Peptidase M84 mediated cellular apoptosis, PA-1 and SKOV3 cells were pre-incubated with either NFκB inhibitor (MG132) or p38 inhibitor (SB203580) or both followed by Peptidase M84 treatment for 18 h. When cells were pre-incubated with MG132 only 58 % of PA-1 and 49 % of SKOV3 cells showed apoptosis. Cells pre-incubated with SB203580 only 51 % of PA-1 and 41 % of SKOV3 cells showed apoptosis. Importantly, the majority of the cells undergoing apoptosis under the aforementioned conditions were mostly in the early apoptotic phase. Interestingly, when both inhibitors were applied together no significant apoptosis was noticed. Our findings suggest activation of NFκB and p38 is essential to trigger apoptosis by Peptidase M84 in ovarian cancer cells. Furthermore, no significant apoptosis was noticed when Peptidase M84 treated PA-1 and SKOV3 cells were pre-incubated with ROS quencher (NAC) for 1 h. This observation suggested the major involvement of ROS in Peptidase M84 induced apoptosis in ovarian cancer cells (Fig. 7C-E).

Taken together our data illustrated that Peptidase M84 induced apoptosis in ovarian cancer cells effectively targeting PAR-1 with the specific involvement of p38, NF-κB and ROS (Supplementary Fig S3 A-D).

We have stated earlier that Peptidase M84 treatment promoted ROS generation in ovarian cancer cells. Interestingly, cells pre-treated with ML-161 displayed no significant ROS generation even after Peptidase M84 treatment for 18 h. In addition to this observation, PAR-1 silenced cells also showed similar results even after Peptidase M84 treatment (Fig. 7F-H). Western blot confirmed the knockdown of PAR-1 by si-RNA (Fig. 7I). Inhibition studies with MG132 and SB203580 further strengthened that blockade of NFκB and p38 impaired further ROS production in ovarian cancer cells (Supplementary Fig S3 B-D). These results indicated that PAR-1 induction by Peptidase M84 promoted ROS generation through activation of NFκB and p38 in ovarian cancer cells.

### 3.9 Peptidase M84 treatment improved the survival of ID8 bearing mice and impeded body weight increase due to ascites accumulation and also affect the viability of ID8 cells *in vivo*

Tumour was induced intraperitoneally with ID8 cells (5X10^6^) and the survival rate and the alterations in body weight were observed in C57BL/6 mice (Fig. 8A). In the ID8 control mice, the rate of survival was 60 % after 30 days and 10 % after 60 days. When 3.0 µg/ml Peptidase M84 was injected at a weekly interval for 7 successive weeks the rate of survival was increased to 90 % after 30 days and 70 % after 60 days (Fig. 8B). The body weight of ID8 control group enhanced from 25 g (0 day) to 45 g (after 60 days of tumour inoculation). On the other hand, in Peptidase M84 treated group and buffer control group no significant alterations in body weight were noticed (Fig. 8C and Supplementary Fig. S4 A, B). Inactivated Peptidase M84 (EDTA treated) failed to ameliorate the survival rate of ID8 induced mice and also showed an increase in body weight (Fig. 8C).

**Fig. 8.**
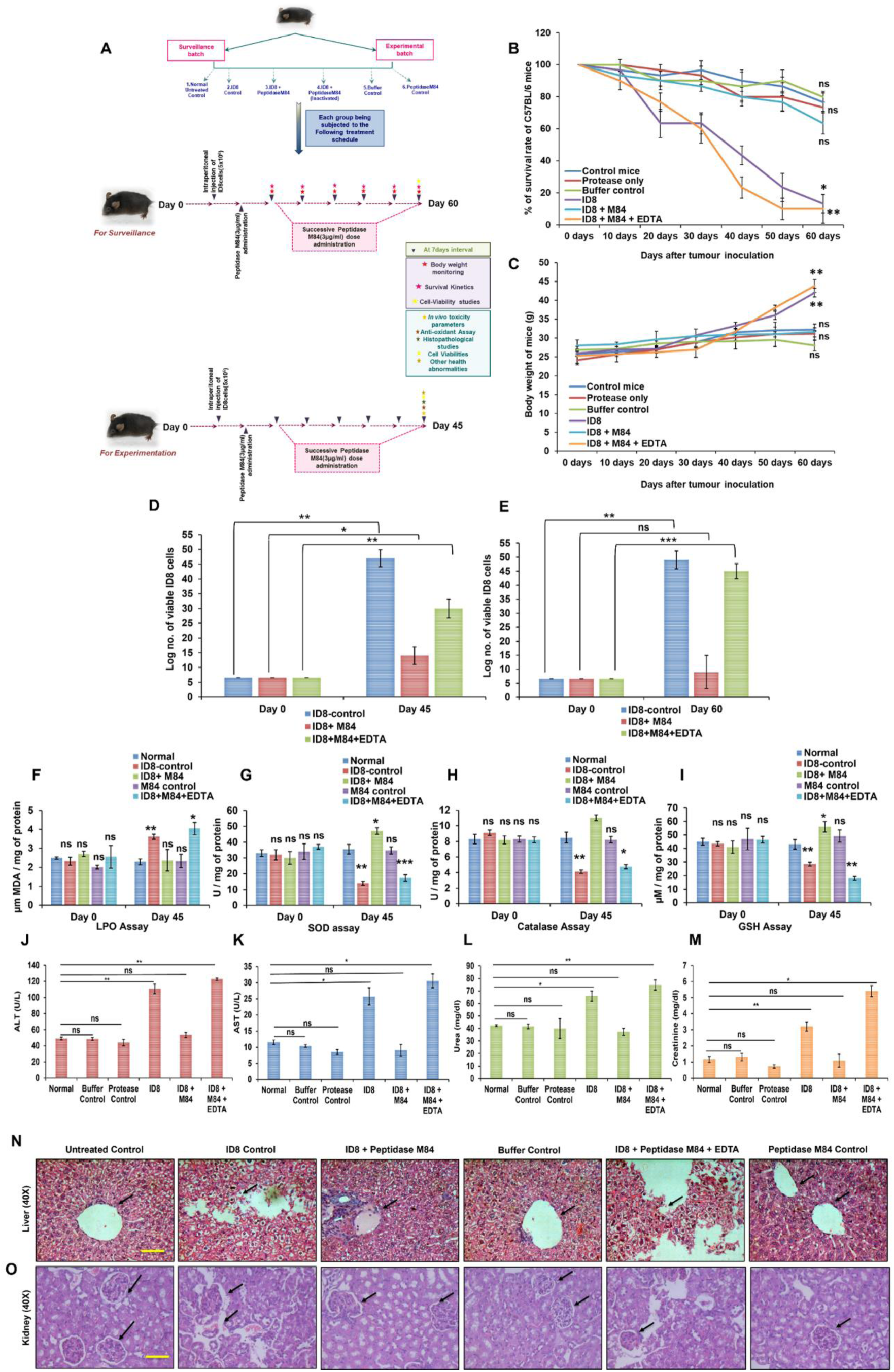
Effect of Peptidase M84 on histological, biochemical parameters, survival rate and body weight of tumour induced mice. **A** Illustration of the *in vivo* study to determine the anticancer potential and non-toxicity of Peptidase M84. **B** Peptidase M84 treatment showed promising improvement in survival rate in ID8 induced intraperitoneal mice model. Peptidase M84 in presence of EDTA showed similar mortality rate as tumour control group. **C** Body weight after Peptidase M84 administration in mice. **D, E** Viable ID8 cells measured at different time points by the trypan blue exclusion method. **F-I** Biochemical parameters of cellular oxidative stress (LPO, SOD, CAT and GSH) were measured in serum samples of mice. Bar diagrams depicted a significant decrease in LPO levels and increase in SOD, GSH and catalase levels due to Peptidase M84 treatment as compared to ID8 control group. **J-M** Liver and kidney functionality test (ALT, AST, Urea and Creatinine) showed the recovery of liver and kidney cells from ID8 mediated damage after Peptidase M84 treatment. **N, O** Histological studies by H&E staining showed that Peptidase M84 treatment improved the damaged architecture of liver and kidney tissue caused by tumour induction. While arrows indicated intact membrane integrity in control and Peptidase M84 treated set and disrupted tissue architectures in ID8 control and Peptidase M84-pre-treated with EDTA set. In each panel, error bars were calculated based on results obtained from a minimum of three independent experiments (n=3). All statistical analysis was done by applying Student’s t-test (two-tailed). Data are expressed in ± SEM. In all panels, ns p>0.05, *p≤0.05, **p≤0.01, and ***p≤0.001.

In addition, viable ID8 cells were evaluated by trypan blue staining. At the inception of this investigation ID8 cells (5X10^6^) were inoculated in the peritoneum cavity of mice and there was a gradual proliferation of viable ID8 cells in tumour control group. Peptidase M84 treatment caused a significant decline in the count of viable ID8 cells after 45^th^ and 60^th^ days. Peptidase M84 inactivated by EDTA did not affect the viability of ID8 cells at all (Fig. 8D, E). Altogether our findings elucidated that the antitumour property of Peptidase M84 was linked to its proteolytic activity.

### 3.10 Peptidase M84 augmented oxidative stress in mice

To evaluate the Peptidase M84 mediated oxidative stress, biochemical tests were performed with different phase II detoxifying enzymes and LPO in the serum samples of mice. The level of LPO in serum is considered a marker for cellular oxidative stress. After 45 days of treatment, the number of viable ID8 cells significantly lessened in group-3 as compared to group-2 and the level of LPO in the serum was also found to be lowered in group-3 than to that of tumour control group-2. This indicated that Peptidase M84 itself could induce oxidative stress. After 45 days of Peptidase M84 treatment, the LPO level in group-3 was at normal levels as compared to group-1. The LPO levels of group-4 were found similar to group-2 after 45 days of treatment (Fig. 8F).

The levels of catalase, SOD and GSH in serum were relatively low because of excessive oxidative stress. After 45 days of Peptidase M84 treatment, the number of viable ID8 cells was significantly declined in group-3 as compared to group-2 and increase in the levels of these enzymes was found in group-3 than to group-2. In addition to the above, the levels of catalase, GSH and SOD in serum of group-4 were noticed as similar as group-2 after 45 days of treatment. Interestingly, there was no significant difference observed in any of the above-mentioned biochemical markers when normal animals were treated with Peptidase M84 (group-6) (Fig. 8G-I). These observations clearly pointed out that Peptidase M84 triggered apoptosis in ID8 cells via oxidative stress mediated damage.

### 3.11 Studies based on the biochemical index of the liver and kidney of mice revealed Peptidase M84 did not cause any significant toxicity

To observe any adverse effects of exposure to Peptidase M84, a chronic toxicity study for 45 days was also been performed on experimental C57BL/6 mice. Alterations in the level of biochemical parameters such as SGOT (AST), SGPT (ALT) (parameters for drug-induced liver injury) or creatinine and urea (parameters for toxin-induced impaired renal function) in the serum sample of experimental mice implicate hepatotoxicity and nephrotoxicity. ALT and AST activity was significantly upregulated and creatinine, urea levels were reduced significantly in the tumour control group when compared to the normal group. However, no such significant difference was noticed in all the above-mentioned indexes in the only protease treated group with respect to normal untreated group. Our results deciphered that both liver and kidney toxicity parameters remained within the normal range after treatment with 3.0 μg/ml (12.0 µg / kg of body weight) of Peptidase M84 for up to 45 days of continual treatment (Fig. 8J-M). No adverse effects were observed concerning normal untreated mice. Upon evaluating the chronic toxicity of Peptidase M84, it did not cause any significant toxicity as all the parameters for liver and kidney toxicity lied within normal limit. We further validated those findings by histopathological analysis.

### 3.12 Peptidase M84 treatment showed recovery of the damaged architecture of liver and kidney tissue of host system and did not cause any significant toxicity

In the current investigation, histopathological examination was executed on sections from kidney and liver tissues of all six groups of experimental animals after 45 days of treatment using haematoxylin-eosin staining to assess whether there was any inconsistency with the biochemical markers. Histopathological sections of normal liver unraveled healthy morphology of hepatocytes; normal kidney tissues displayed healthy glomeruli, vessels and tubules. However, tumour control group (group-2); showed disrupted architecture with moderate to dense inflammation with damaged hepatocytes in liver, and extensively damaged glomerular structure in the kidney. After Peptidase M84 treatment (group-3) this damage was observed to be recovered and healthy tissue structure prevailed in liver and kidney tissue. In comparison with the untreated control group, no noticeable anomalies in the histopathology of the kidney, and liver of the Peptidase M84 treated group-3 and group-4 were detected, following 45 days of continual treatment. Morphology of the liver of only Peptidase M84 treated mice was almost similar to the untreated group. Morphology of the livers from group-6 showed no significant change from that of the untreated control. As observed in untreated control, in addition to the radial distribution pattern of hepatocytes encircling the central vein was obvious in the protease control and vehicle control group after treatment. Radially arranged hepatocytes (HC) lacking Kupffer cells (KC) and fine integrity in the liver tissue, depicting the absence of any chronic toxicity (Fig. 8N).

Notable differences after 45 days of constant treatment with Peptidase M84 between renal corpuscles (RC) and glomerular tufts of kidney tissue were not found when compared to the untreated group. Moreover, paucity of tubular dilation, glomerular infiltration and necrosis indicated the non-existence of acute inflammation (Fig. 8o). Hence, this implies that 3.0 µg/ml (12.0 µg/kg of body weight) of Peptidase M84 for the mentioned period is significantly non-toxic, non-hazardous to normal tissue of mice and does not affect the survival of mice (Fig: 8N, O).

## 4. Discussion

Emerging complexities and side effects associated with conventional chemotherapy necessitate the quest for new therapeutic agents for ovarian cancer treatment. In this study, we designed a novel approach to isolate and characterize proteases of environmental microbial origin for apoptotic properties to curb the growth of ovarian cancer cells. For this purpose, we assessed over 200 environmental isolates to finally select a protease secreting strain ‘GDL-186’ which was identified as *Bacillus altitudinis.* Reported to be first isolated from air samples of high altitudes in India, *Bacillus altitudinis* was found to secrete extracellular proteases potentiating apoptosis in ovarian cancer cells in our study [66]. Popularly, *Bacillus* spp. is widely studied for its extracellular alkaline protease secreting properties which also make it industrially important [67]. In our attempt to find a protease with pro-apoptotic properties, we identified Peptidase M84 which is a key secretory metallo-protease of *Bacillus altitudinis* [68]. A recent report revealed the anti-proliferative activity of the organic extract of *Bacillus altitudinis* MTCC13046 against human hepatocellular carcinoma cells [69]. We obtained two major bands at 25kDa and 16 kDa in SDS-PAGE for this purified protease which was homologous to the amino acid sequences of Peptidase M84 from *Bacillus altitudinis* in Uniprot and NCBI databases. This protease is comprised of a conserved metallo-protease domain with three histidine residues in its active site, which is also reported to be present in metzincin metallo-protease from *Bacillus intermedius* [60]. The optimal protease activity of Peptidase M84 was observed in normal physiological pH (7.0–9.0) and temperature (37°C-40°C) that got inhibited by EDTA and 1,10 phenanthroline. This aided us to characterise Peptidase M84 as a Zn^++^-dependent metalloprotease.

To our knowledge, this is the first report to purify and characterise Peptidase M84 from an environmental isolate of *Bacillus altitudinis.* Previously, a polypeptide PBN118 isolated from a marine Bacillus was reported to be similar to the Peptidase M84 from *Bacillus pumilus.* PBN118 prevented migration and suppressed invasion of human hepatocellular carcinoma cells [70]. Peptidase M84, here also triggered apoptosis in both human and mouse ovarian adenocarcinoma cells in a dose-dependent manner. Moreover, this pro-apoptotic response of Peptidase M84 was inhibited by EDTA treatment which helped us confirm that its apoptotic response was a result of its proteolytic activity. Interestingly, Peptidase M84 did not show any significant effect on IOSE and PEMФ cells which were the normal cells used in this study. Additionally, Peptidase M84 turned out to be anti-proliferative in nature when it reduced the expression levels of Ki-67, a proliferative antigen in PA-1 and SKOV3 cells. Based on these findings, we were intrigued to decipher the detailed molecular mechanisms underlying the Peptidase M84 induced apoptosis in ovarian cancer cells. Since more than 90% of all ovarian malignancies are classified as EOC including the aggressive high-grade serous carcinoma category [1], we considered that initial testing of Peptidase M84 for its apoptosis-inducing capabilities in *in vitro* setup could enable the designing of a better treatment rationale.

The canonical intrinsic pathway of apoptosis depends on the delicate balance between pro-apoptotic Bax and anti-apoptotic Bcl2 [71]. We observed a significant increase in Bax to Bcl2 ratio, and a rise in cytosolic mitochondrial cytochrome c concentrations in addition to caspase 9 and 3 activation following Peptidase M84 treatment in SKOV3 as well as PA-1 cells. These observations delineated an induction of intrinsic apoptosis pathway in these malignant ovarian cells by Peptidase M84 specifically. Enhanced oxidative stress owing to the production of higher ROS and impaired redox balance is intrinsic to the sustenance of any neoplastic cells unlike a normal healthy cell [25, 26]. In coherence with this fact, we also noted that Peptidase M84 abnormally increased intracellular ROS levels in PA1, SKOV3 and ID8 cells in a time dependent manner which resulted in their apoptotic death. Contrastingly, Peptidase M84 failed to impart any such change in normal IOSE cells where ROS was quantitated to be very less with respect to the *in vitro* malignant setups. This observation was further confirmed by the inhibition of ROS by pre-incubation with NAC (a cellular ROS quencher) before treatment with Peptidase M84 in ovarian cancer cells. Overproduction and accumulation of ROS is known to modify nucleotide, break DNA strand and facilitate chromosomal rearrangements, contributing to the deregulation of a wide range of signalling pathways [72]. Peptidase M84 treatment induced ROS mediated DNA damage in ovarian cancer cells which intrigued us to check the status of mitochondrial membrane potential by JC-1 staining as ROS overproduction is always positively associated with mitochondrial membrane depolarization and reduced cell viability. A disruption in mitochondrial membrane potential by increased cellular ROS was apparent in JC1 stained ovarian cancer cells. These findings aligned with a rise of cytochrome c as detected in the cytosolic fractions of Peptidase M84 treated ovarian cancer cells which might have leaked out of a depolarized mitochondrial membrane to trigger apoptosis cascades involving caspases, apoptosome complex and PARP [65, 73]. Enzymatic activity of PARP is found to be increased in the cells under stressed conditions. Intact PARP expression is associated with DNA repair system which in turn saves the cancer cells from apoptotic cell death. The presence of cleaved PARP and a decrease in Bcl-2 muddle the apoptotic balance and drive the cell towards the gateway of apoptosis, as it restricts cell repair [74]. We evidenced Peptidase M84 to induce PARP cleavage in ovarian cancer cells which corroborated with an increased DNA damage. This PARP activating feature of Peptidase M84 is promising enough since FDA has approved Olaparib (AZD2281) which is a PARP inhibitor for conventional ovarian cancer treatment [75].

The major signalling pathways of proteases are initiated by the cleavage of PARs which play an important role both in cancer progression and apoptosis of malignant cells in a cue-based manner [28]. PAR1 was reported to be overexpressed in explants of human ovarian cancer tissues compared to normal ovarian cells [29, 34]. Gingipain-R (RgpB) a cysteine protease isolated from *Porphyromonas gingivalis* was the earliest reported microbial protease that acted by cleaving a model peptide representing a cleavage site of PAR-2 [76]. RgpB has been also reported to promote platelet aggregation through the activation of PAR-4 and PAR-1 [76]. We noted an overexpression of PAR1 at both transcriptional and translational levels in Peptidase M84 treated ovarian cancer cells compared to their untreated counterparts. Interestingly, no significant alteration in the expression level of PAR1 in Peptidase M84 treated normal ovarian epithelial cells was seen. Several evidences identified PAR-1 as a tumour promoter as it was associated with the induction of angiogenesis, invasion and also metastasis in ovarian, breast adenocarcinoma cells along with xenograft models [77, 78]. However, thrombin is found to mediate apoptosis in a dose-dependent way through modulation of PAR-1 in tumour cells [33]. Previously we reported that metallo-protease HAP from *V. cholerae* could cause overexpression of PAR-1 by inducing its cleavage [37]. Nevertheless, such a paradox is not limited only to PAR-1 only. Biomolecules like granulocyte/macrophage colony-stimulating factor, RAR-β2, E-cadherin, CD44, α/β-catenin and CAV1 have also been reported to impart such contrasting functions in tumourigenesis [79, 80]. In order to reveal the relationship between alteration in PAR-1 expression and apoptosis in the presence of Peptidase M84 in ovarian cancer cells, we performed gene knockdown, signalling inhibition and immunoprecipitation studies. Our findings also demonstrated that Peptidase M84 could specifically bind and interact with PAR-1. Moreover, Peptidase M84 failed to induce apoptosis in PAR1 silenced ovarian cancer cells. Furthermore, ML161, a PAR1 mediated Gq signalling inhibitor [81] also showed a significant decrease in apoptosis in Peptidase M84 treated cells. This suggests that Peptidase M84 caused PAR-1 activation through Gq signalling induction. Cumulative data strengthened the possible correlation regarding the role of PAR-1 in induction of apoptosis and established PAR-1 as a novel oncogenic target for Peptidase M84 mediated apoptosis in ovarian cancer cells. PAR1 has emerged as a promising hot spot target for chemotherapeutic drugs in recent years as well. PAR1 targeting drugs like vorapaxar and atropaxar have entered phases of clinical trials [31].

Thrombin-mediated PAR-1 activation regulates major signalling pathways such as p38 MAPK [82], and NFκB [83]. Herein, inhibition studies using NFκB and p38 inhibitors confirmed the involvement of NFκB and MAPK in Peptidase M84 mediated apoptosis in PA-1 and SKOV3 cells. Pre-treatment with NFκB and p38 inhibitors downregulated cellular apoptosis even after Peptidase M84 treatment compared to the absence of any prior inhibition. Interestingly, besides the upregulation of the phosphorylated form of p38, a downregulation of p-ERK1/2 level was observed in PA-1 and SKOV3 cells upon Peptidase M84 treatment. Blockade of AKT/ERK pathway and enhanced phosphorylation levels of p38, contributing majorly in apoptosis in response to different stimuli in cancer cells [84]. NF-κB regulates the expression of genes in cellular processes like cellular proliferation, stress responses and apoptosis under different stimulations [85]. Evidences showed that p38 and NFκB can augment the induction of cellular ROS levels [86]. Models representing oxidative stress have been explored to understand the effects of oxidative stress on NF-κB related activities. Besides, ROS can also regulate either activation or repression of NF-κB signalling depending on phase and context [87, 88]. Our earlier studies on PAR-1 mediated apoptosis in colon carcinoma showed ROS generation in cancer cells by means of activation of NF-κB and MAPK by HAP from *V.cholerae* and inhibition of PAR-1 impeded ROS generation [37]. Here, we found inhibition of apoptosis when Peptidase M84 treated cells were pre-incubated with NAC and partial inhibition after individual application of NFκB and p38 inhibitors. However, there was increase in ROS generation in ovarian cancer cells as an earlier response (6 h) to Peptidase M84 treatment which increased significantly in a time dependent manner. In light of these findings, we tried to establish the relationship between activation of NFκB and oxidative stress induced by Peptidase M84 in ovarian cancer cells. Enhanced ROS levels promoted the activation of NFκB and simultaneously triggered ROS generation further. Phosphorylation of p38, a major component of MAPK pathway is linked with ROS generation and cell proliferation in the majority of the events [86, 89]. However, when p-p38 levels cross the critical threshold in conjugation with the downregulation of ERK, cells get immensely stressed to meet death [90]. Our findings also revealed inhibition of ROS production upon blockade of PAR-1 mediated signalling by Si-RNA and ML-161 in ovarian cancer cells. In addition, there was no significant expression of PAR-1 in IOSE cells. Hence, the load of cellular ROS level was found to be lower in IOSE cells even after Peptidase M84 treatment as compared to cancer cells, as there was no effective alteration in PAR-1 activity or expression. Thus, Peptidase M84 failed to trigger the intrinsic apoptotic pathway in normal IOSE cells.

These results indicated that NFκB, p38, ERK1/2 and cellular ROS level regulate Peptidase M84 induced apoptosis by playing decisive roles either individually or perhaps in a collaborative manner in ovarian cancer cells.

Finally, to validate our findings *in vivo* we investigated the role of Peptidase M84 in a syngeneic mouse model. ID8 is a spontaneous murine ovarian adenocarcinoma cell line used to study ovarian tumour biology. This cell line is advantageous due to its efficacy to develop tumours in both solid and ascitic forms [53, 54]. Herein, we report that Peptidase M84 treatment in ID8 mice resulted in significant inhibition body weight increase due to less ascites formation. An increase in the survival rate of the ID8 bearing mice following Peptidase M84 treatment as compared to the tumour control group indicated that Peptidase M84 treatment affected the viability of cancer cells *in vivo*. However, Peptidase M84 pre-treated with EDTA was found to be ineffectual and did not show any kind of alterations. This indicated the antitumour activity of Peptidase M84 was linked to its proteolytic activity like Serratia protease or HAP. In the ID8 bearing mice, the cellular ROS level was increased significantly after Peptidase M84 treatment. Increase in the Peptidase M84 induced oxidative stress corroborated with a rise in the levels of GSH, SOD and catalase along with a reduced LPO titres in the ID8 bearing mice receiving Peptidase M84 treatment as compared to the ID8 control mice. Moreover, the reduction in viable ID8 cell count in mouse peritoneum indicated that Peptidase M84 treatment decreased the survivability of tumour cells and strengthened the remedial effect chemotherapy. The histopathological studies illustrated that the cellular architecture and morphology of the liver and kidney tissues of mice were not affected by Peptidase M84. Hence, it is significantly non-toxic for normal tissue of mice. However, Peptidase M84 treatment was capable of recovering the damaged tissue morphology in ID8 bearing mice. The increase in the levels of SGOT, SGPT, urea and creatinine in ID8 control group with respect to the group of mice treated with Peptidase M84 in regular intervals further validated our findings. No significant mortality or toxic symptoms were noticed in only Protease control group. Thus, our findings demonstrated a reduction in cancer cell proliferation via oxidative stress mediated apoptosis without causing harm to normal healthy cells following Peptidase M84 treatment *in vivo*.

## 5. Conclusion

Collectively our findings demonstrated the role of Peptidase M84 as a novel anticancer molecule that induces ROS mediated apoptosis via effective interaction with PAR-1, ensuing prevalent signalling changes that eventually elicit anticancer effects in ovarian cancer cells. Our study envisaged an activation and overexpression of PAR-1 by Peptidase M84 induced enhancement of cellular ROS and activation of NFκB, p38 followed by downregulation of pERK1/2. ROS mediated DNA damage induced changes in mitochondrial membrane potential and triggered the release of mitochondrial cytochrome c which subsequently activated caspase 3 ultimately promoting Bax, Bcl2 mediated intrinsic pathway of apoptosis. Enhanced ROS level in malignant cells allowed to meet the threshold level of cellular ROS that affect cell survival earlier than the normal healthy cells ultimately triggering apoptotic cell death. We also report that Peptidase M84 triggered apoptosis in *in vivo* mice models without imparting any effect on human and mouse normal cells in addition to the liver and kidney tissue of C57BL/6 mice. Peptidase M84 also increased the lifespan of tumour bearing mice. Thus, Peptidase M84 warrants further exploration as a new antitumour agent. This may raise awareness among the researchers to explore other protease producing microbes along with their specific molecular targets to cure cancer.

## 6. Limitations of the study

Unlike thrombin, the Peptidase M84 mediated exact PAR-1 cleavage site is in the process of identification which will be the major focus of our future study. Further investigations may be executed in other experimental models and clinical settings to enlist Peptidase M84 as an anticancer drug, which may widen the amplitude for a long-lasting targeted therapy for ovarian cancer.

## Acknowledgements

Mr. Niraj Nag is thankful to University Grants Commission, India for providing fellowship under the prestigious UGC-NET-JRF fellowship award. We wish to express our gratefulness to Dr. Shanta Dutta (Director, ICMR-NICED, India) for providing us with the central research instrumentation facility of NICED. Mr. Biplab Roy (ICMR-NICED, India) is gratefully acknowledged for helping us in the screening of environmental isolates and animal experiments. We thankfully acknowledge Mr. Mainak Chakraborty (ICMR-NICED, India), Dr. Moumita Bhaumik (ICMR-NICED, India), Dr. Atish Barua (Tufts University, USA), Mr. Prolay Halder (ICMR-NICED, India), Dr. Hemanta Koley (ICMR-NICED, India), Dr. Sutapa Mukherjee (CNCI, India) Mr. Nanda Singh (ICMR-NICED, India) for their technical support and co-operation during the study.

## Author’s contribution

**Niraj Nag** conceptualised the study, investigated, performed experiments, wrote the original draft, reviewed, data curation, analysis and validation of the data; **Tanusree Ray** performed experiments, reviewed; **Rima Tapader** formal analysis, reviewed; **Animesh Gope** performed experiments; **Rajdeep Das** analysis of confocal imaging data; **Elizabeth Mahapatra** formal analysis, reviewed; **Saibal Saha** formal analysis; **Ananda Pal** flowcytometric data acquisition and formal analysis; **Parash Prasad** provided resources; **Shruti Chatterjee** provided resources; **Sib Sankar Roy** provided resources; **Amit Pal** conceptualised the study, supervised the research work, funding acquisition, data curation, reviewed and approved the final manuscript.

## Ethical approval for animal studies

The experimental design of the present study involving animals was approved by the Institutional Animal Ethics Committee (IAEC) (License No: PRO/168/Jan 2022), NICED, Kolkata, India. The central experimental animal facility provided the mice with sterile paddy husk bedding encased in pathogen-free polypropylene cages. The temperature and humidity were kept constant (23 ± 2°C and 55 ± 10 %) with alternate light and dark cycles (12 h/12 h). Animals were fed with a standard food pellet diet (EPIC rat and mice pellet from Kalyani Feed Milling Plant, Kalyani, West Bengal, India) and drinking water *ad libitum*. In each experiment, the mice were acclimatised for three days before being divided into distinct subgroups. The IAEC is formed as per the guidelines of the Committee for the Purpose of Control and Supervision of Experiments on Animals (CPCSEA), Ministry of Environment, Forests and Climate Change, Government of India. For the experimental purpose, euthanasia was done as per CPCSEA guidelines

## Availability of data

The datasets generated during and/or analysed during the current study are available from the corresponding author on reasonable request.

## Consent to publish

Not applicable.

