## Supplementary Figures for "Metallo-protease Peptidase M84 from *Bacillus altitudinis* induces ROS dependent apoptosis in ovarian cancer cells by targeting PAR-1"

### Title Page

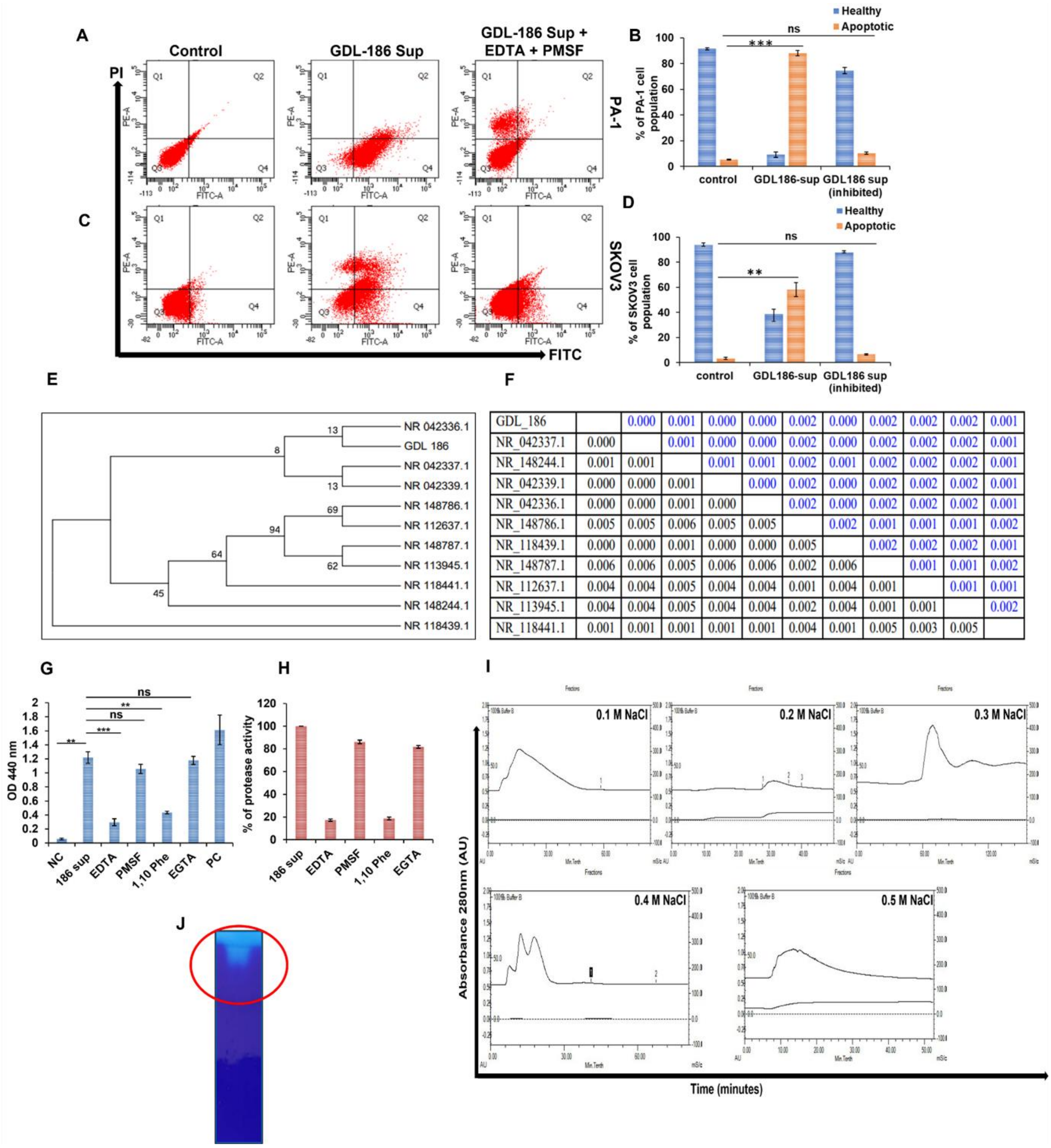


| | | | | | | | | | | | |
|---|---|---|---|---|---|---|---|---|---|---|---|
| GDL_186 | | 0.000 | 0.001 | 0.000 | 0.000 | 0.002 | 0.000 | 0.002 | 0.002 | 0.002 | 0.001 |
| NR_042337.1 | 0.000 | | 0.001 | 0.000 | 0.000 | 0.002 | 0.000 | 0.002 | 0.002 | 0.002 | 0.001 |
| NR_148244.1 | 0.001 | 0.001 | | 0.001 | 0.001 | 0.002 | 0.001 | 0.002 | 0.002 | 0.002 | 0.001 |
| NR_042339.1 | 0.000 | 0.000 | 0.001 | | 0.000 | 0.002 | 0.000 | 0.002 | 0.002 | 0.002 | 0.001 |
| NR_042336.1 | 0.000 | 0.000 | 0.001 | 0.000 | | 0.002 | 0.000 | 0.002 | 0.002 | 0.002 | 0.001 |
| NR_148786.1 | 0.005 | 0.005 | 0.006 | 0.005 | 0.005 | | 0.002 | 0.001 | 0.001 | 0.001 | 0.002 |
| NR_118439.1 | 0.000 | 0.000 | 0.001 | 0.000 | 0.000 | 0.005 | | 0.002 | 0.002 | 0.002 | 0.001 |
| NR_148787.1 | 0.006 | 0.006 | 0.005 | 0.006 | 0.006 | 0.002 | 0.006 | | 0.001 | 0.001 | 0.002 |
| NR_112637.1 | 0.004 | 0.004 | 0.005 | 0.004 | 0.004 | 0.001 | 0.004 | 0.001 | | 0.001 | 0.001 |
| NR_113945.1 | 0.004 | 0.004 | 0.005 | 0.004 | 0.004 | 0.002 | 0.004 | 0.001 | 0.001 | | 0.002 |
| NR_118441.1 | 0.001 | 0.001 | 0.001 | 0.001 | 0.001 | 0.004 | 0.001 | 0.005 | 0.003 | 0.005 | |

**Supplementary Fig. S1 A, C** Flow cytometric studies to detect apoptosis with overnight grown culture supernatant from isolate GDL-186 in PA-1 and SKOV3 cells indicated the presence of a microbial extracellular protease with apoptosis inducing capabilities. Overnight grown culture supernatant of isolate GDL-186 priorly incubated with EDTA and PMSF failed to induce apoptosis in PA-1 and SKOV3 cells. **B, D** These results are graphically represented. **E** Molecular phylogenetic analysis and the evolutionary history were inferred by using the maximum likelihood method based on the Kimura 2 parameter model. The 'NR' number stands for the respective NCBI accession numbers. **F** Estimates of evolutionary divergence between sequences. The number of base substitutions per site from between sequences is shown. Standard error estimate(s) are shown above the diagonal. Analyses were conducted using the Kimura 2-parameter model. **G, H** Azocasein assay showed protease activity in culture supernatant of *Bacillus altitudinis* strain GDL-186 which showed reduction upon treatment with EDTA and 1,10- phenanthroline but not by PMSF and EGTA. Here culture supernatant from *Vibrio cholerae* C6709 was used as positive control (PC) and nutrient broth was used as negative control (NC). **I** Chromatogram profile of DEAE-52 ion-exchange chromatogram profile of binding fractions of the crude protein extract from the culture supernatant of *Bacillus altitudinis* strain GDL-186 eluted by gradient of NaCl (0.1 M – 0.5 M). **J** Native gelatin zymogram profile of purified Peptidase M84 showed proteolytic degradation of gelatine. All statistical analysis was done by applying Student's t-test (two tailed). Data are expressed in ± SEM. In all panels, ns $p>0.05$, *$p\leq0.05$, **$p\leq0.01$, and ***$p\leq0.001$. In each panel, error bars were calculated based on results obtained from minimum of three independent experiments.

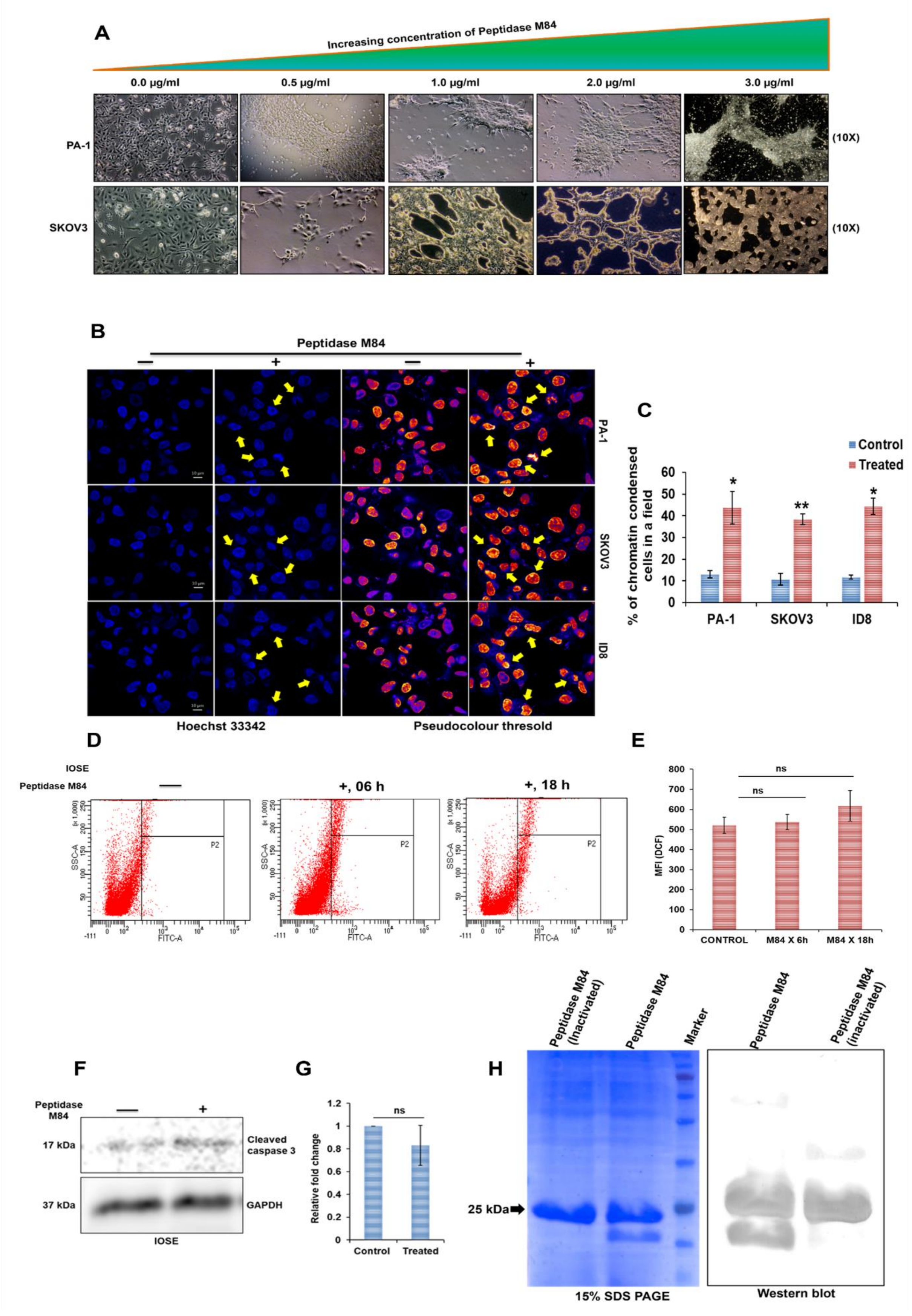


**Supplementary Fig. S2 A** PA-1 and SKOV3 cells grown in tissue culture dishes were treated with different concentrations of Peptidase M84 for 18 h. Cells were observed with a phase-contrast microscope. Cell distending and rounding effects of Peptidase M84 were noticed in lower and higher concentrations respectively**.** **B** PA-1, SKOV3 and ID8 cells were stained with Hoechst 33342 after treatment with Peptidase M84. The representative images of Hoechst 33342 stained nuclei (arrow) are shown. **C** Percentage of condensed nuclei is represented graphically. **D** Detection of ROS generation in IOSE cells after Peptidase M84 treatment by DCFDA staining. No significant ROS generation was found after 18 h of treatment. **E** Bar diagram of mean fluorescence intensity of DCF is represented. **F, G** Western blot showed no significant expression of cleaved caspase 3 in Peptidase M84 treated IOSE cells. **H** Detection of two major bands of purified Peptidase M84 from *Bacillus altitudinis* strain GDL-186 by western blot analysis using anti-Peptidase M84 antisera raised in rabbit. All statistical analysis was done by applying Student's t-test (two tailed). Data are expressed in ± SEM. In all panels, ns $p>0.05$, *$p\leq0.05$, **$p\leq0.01$, and ***$p\leq0.001$. In each panel, error bars were calculated based on results obtained from minimum of three independent experiments. Scale bars: 10 μm.

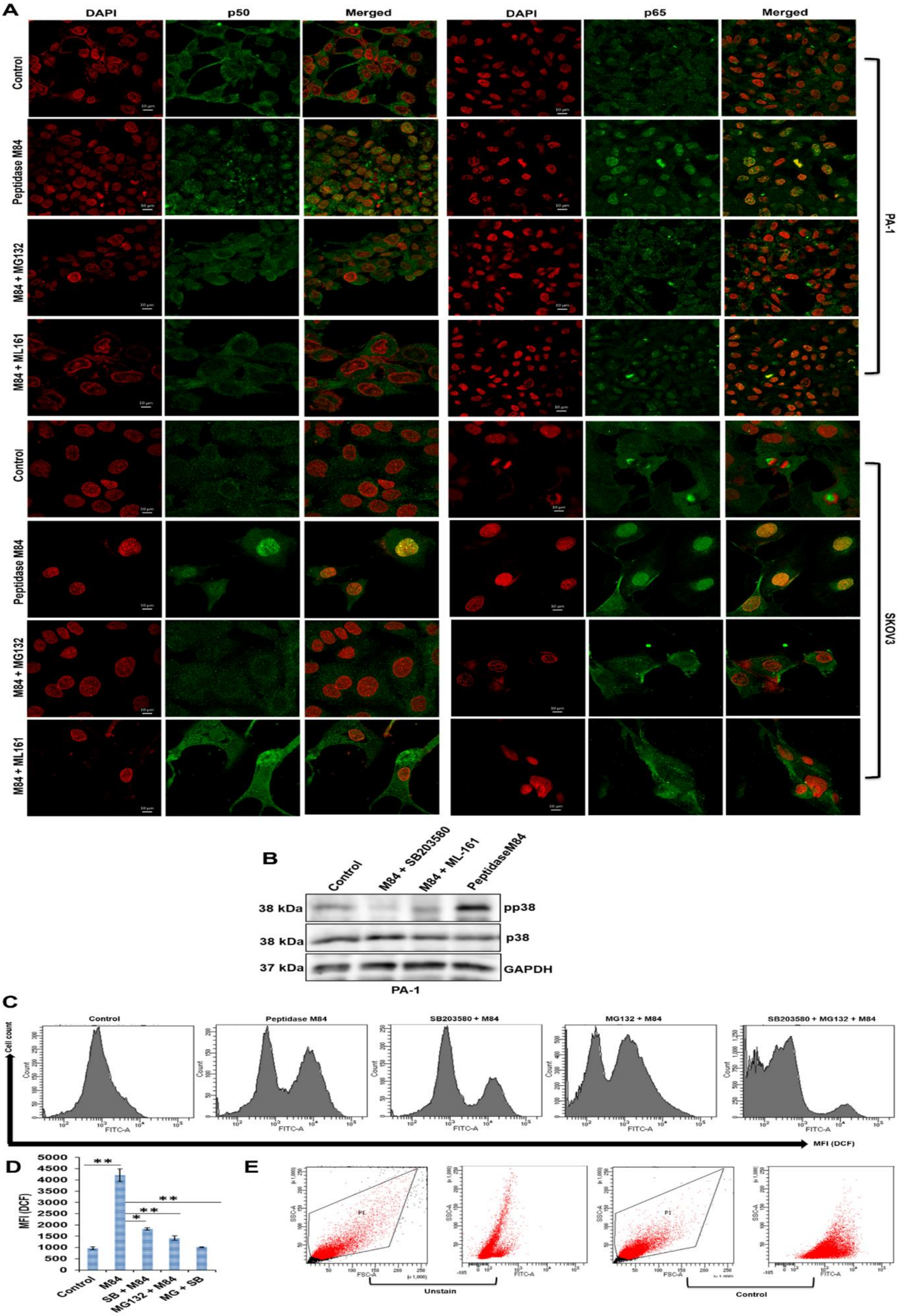

A
DAPI
p50
Merged
DAPI
p65
Merged
Control
Peptidase M84
M84 + MG132
M84 + ML161
PA-1
Control
Peptidase M84
M84 + MG132
M84 + ML161
SKOV3
B
Control
M84 + SB203580
M84 + ML-161
PeptidaseM84
38 kDa
pp38
38 kDa
p38
37 kDa
GAPDH
PA-1
C
Control
Peptidase M84
SB203580 + M84
MG132 + M84
SB203580 + MG132 + M84
Cell count
FITC-A
MFI (DCF)
D
MFI (DCF)
Control
M84
SB + M84
MG132 + M84
MG + SB
E
SSC-A
FSC-A
FITC-A
Unstain
Control

**Supplementary Fig. S3 Signalling Inhibitions to understand Peptidase M84 mediated signalling pathway with MG132 (NFκB inhibitor), SB203580 (p38 inhibitor) and ML161 (PAR-1 inhibitor) A** Inhibition of Peptidase M84 mediated activation and nuclear translocation of NFκB (p50 and p65) by MG132 and ML161 in PA-1 and SKOV3 cells. **B** Pre-treatment of SB203580 and ML161 effectively inhibited the Peptidase M84 activation of p38 in PA-1. **C** Pre-treatment of MG132, SB203580 individually in PA-1 cells partially inhibited ROS generation while pre-treatment of both the inhibitors completely blocked ROS generation. **D** Bar diagram represents the MFI of DCF in PA-1 cells. **E** Dot plot of flow cytometry of the unstained and control samples of PA-1 cells for the same experiment. Scale bars: 10 μM.

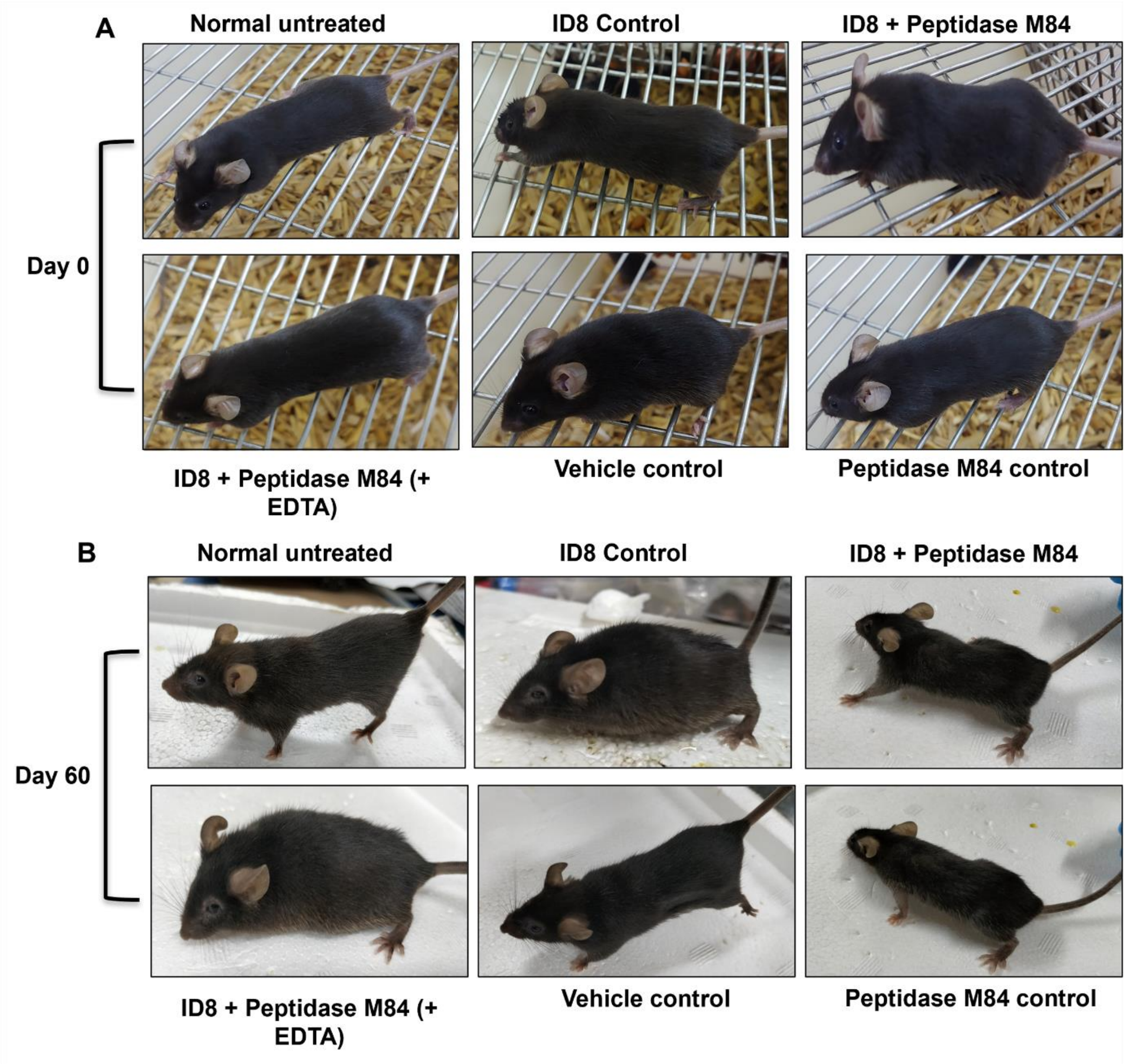


**Supplementary Fig. S4** Effect of Peptidase M84 on ID8 induced intraperitoneal mice model. **A, B** Increase in body weight was observed in ID8 control and Inactivated Peptidase M84 treated ID8 bearing group after 60 days of tumour inoculation. Untreated control group, buffer control group, Peptidase M84 control group and ID8 bearing mice treated with Peptidase M84 group showed no such increase in body weight after 60 days of tumour cell inoculation.
